# Cis-Regulatory Evolution of CCNB1IP1 Driving Gradual Increase of Cortical Size and Folding in primates

**DOI:** 10.1101/2024.12.08.627376

**Authors:** Ting Hu, Yifan Kong, Yulian Tan, Pengcheng Ma, Jianhong Wang, Xuelian Sun, Kun Xiang, Bingyu Mao, Qingfeng Wu, Soojin V. Yi, Lei Shi

**Author notes:** These authors contributed equally in this work (Ting Hu, Yifan Kong, Yulian Tan, Pengcheng Ma).

## Abstract

Neocortex expansion has a concerted relationship with folding, underlying evolution of human cognitive functions. However, molecular mechanisms underlying this significant evolutionary process remains unknown. Here, using tree shrew as an outgroup of primates, we identify a new regulator *CCNB1IP1,* which acquired its expression before the emergence of primates. Following the evolution of cis-regulatory elements, the CCNB1IP1 expression has steadily increased over the course of primate brain evolution, mirroring the gradual increase of neocortex. Mechanistically, we elucidated that CCNB1IP1 expression can cause an increase in neural progenitors through shortening G1 phase. Consistently, the CCNB1IP1 knock-in mouse model exhibited traits associated with enhanced learning and memory abilities. Together, our study reveals how changes in *CCNB1IP1* expression may have contributed to the gradual evolution in primate brain.

## Introduction

Neocortex expansion and intense cortical folding are defining characteristics of the human brain, underlying increased cognitive processing and functions such as human language abilities (*1*). To date, several genes specific to the human lineage have been implicated in the dramatic expansion of human neocortex (*2–8*). For example, human-specific genes *NOTCH2NL* and *ARGAP11B* are preferentially expressed in neural progenitor cells (NPCs), which can promote neurogenesis to enlarge brain size during human evolution (*2, 3, 5*). However, beyond brain size, and the degree of cortical folding measured by the gyrification index (GI), has gradually increased along primate evolution (*1, 9–11*) (**Fig. 1A-B**, *R* = 0.90 versus 0.97). Therefore, in addition to the impacts of human-specific genes, common molecular mechanisms underlying primate neocortex expansion and cortical folding exist.

**Fig. 1.**
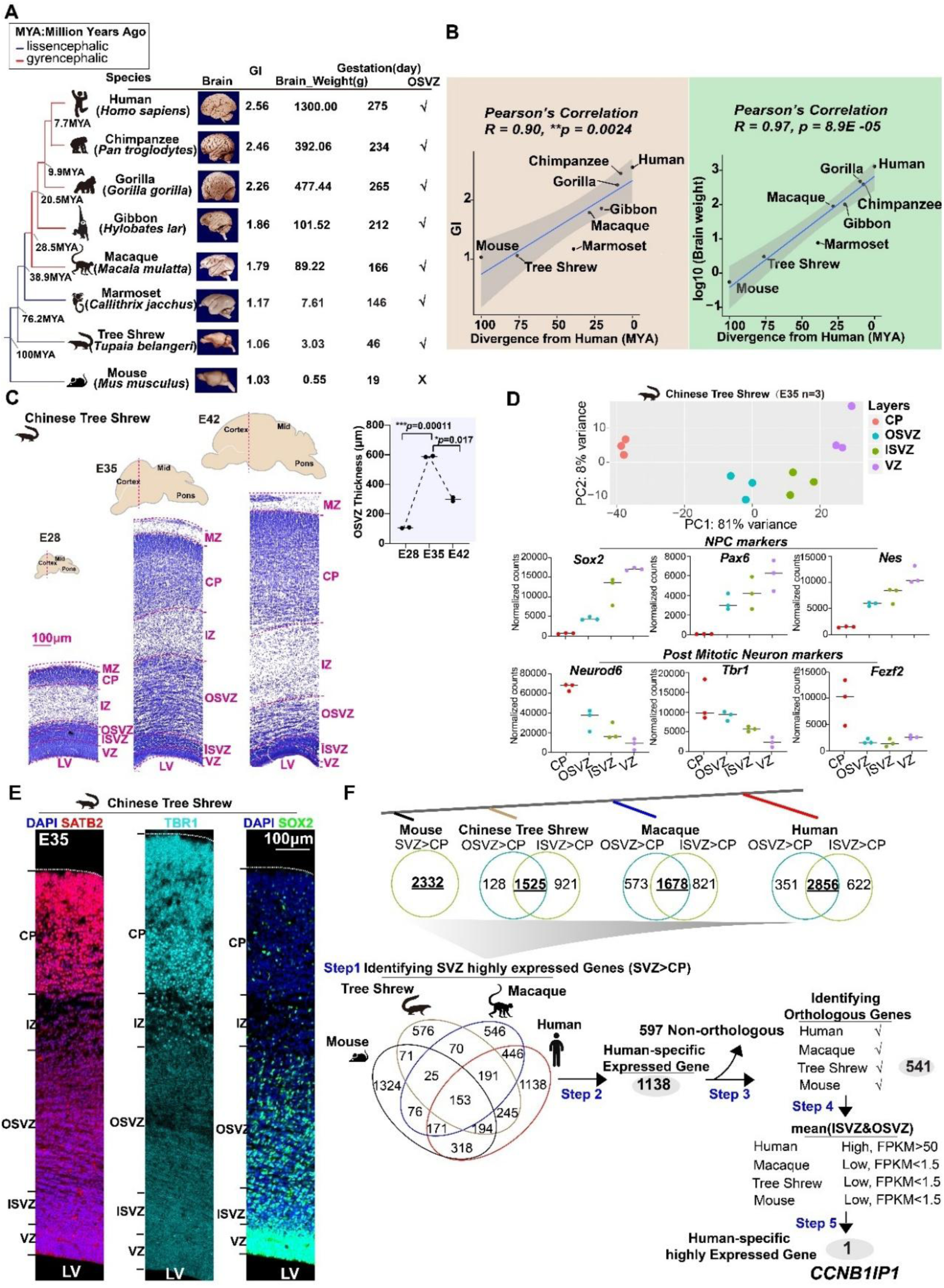
Identification of human specific highly expressed gene *CCNB1IP1*. (**A**) Schematic map showing the evolution of cortical folding and size in primates. Brain images were downloaded from http://brainmuseum.org/. (**B**) (Top) Pearson’s correlation analysis between the gyrification index (GI) and evolutionary divergence (millions of years ago [MYA]). (Bottom) Pearson’s correlation analysis between log transformed brain weight and evolutionary divergence. (**C**) Nissl staining of E28, E35, and E42 Chinese tree shrew cortices and quantification of the OSVZ thickness during tree shrew brain development. * Each point represents a sample. (**D**) PCA map of the frontal cortex layers of Chinese tree shrews at E35 based on the expression levels of all genes. Bottom panel: cell marker analysis with *Sox2*, *Pax6*, and *Nes* for NPCs and Neurond6, Tbr1, and Fezf2 for neurons. (**E**) Immunofluorescence staining of the cortices for the upper layer neuron, deeper layer neuron, and AP markers *SATB2*, *TBR1*, and *SOX2*, respectively, at E35 in Chinese tree shrews. (**F**) Computational pipeline used to identify highly expressed human *CCNB1IP1*.

Cortical folding is developmentally and mechanistically linked to neocortex expansion (*12–14*) which ultimately depends on the activities of neural progenitor cells (NPCs) in the germinal zone (GZ) (*15, 16*). The primary GZ in lissencephalic species (*e.g.* mouse) is the ventricular zone (VZ), which is enriched with apical progenitors (APs) also named as apical radial glial cells (aRGC). In contrast, the primary GZ in gyrencephalic species (*e.g.* human) is the sub-ventricular zone (SVZ), which is enriched with basal progenitors (BPs) (*17*). Intriguingly, the thickness of the SVZ expanded significantly during primate brain evolution, creating an inner (ISVZ) and outer SVZ (OSVZ) (*18, 19*). This expansion is known to be crucial for cortical folding (*15, 20–22*). BPs are generated from APs and consist of intermediate progenitor cells (IPCs) and basal radial glial cells (bRGC). The presence of bRGC is strongly correlated with the degree of gyrification, and bRGCs are rarely found in lissencephalic species (*18, 23*).

Compared with mouse BPs, primate BPs have enhanced self-amplification and self-renewal activities that are closely related to cell cycle length (*24*). For example, primate BPs have a shorter G1 phase than mice, which contributes to the enhanced proliferation of BPs and increased thickness of the SVZ (*25, 26*). In addition, the duration of the cell cycle at mid-neurogenesis is longer in primates than in mice (*26*). The metaphase length of progenitors in humans is approximately 40%–60% longer than that in great apes (*27*). These observations underscore the multifaceted developmental and molecular trajectories underlying human and nonhuman primate brain evolution.

To further our understanding of the molecular mechanisms underlying cortical evolution and complement studies using primate fetal development (*2, 3, 5, 28–32*), we used the Chinese tree shrew (*Tupaia belangeri chinensis*), which is a more closely related to primates than rodents (*33–36*). Importantly, compared with mice, Chinese tree shrews have a significantly expanded SVZ that is particularly enriched in BPs (*37, 38*), indicating that the expansion of SVZ occurred before the emergence of primates. Intriguingly, tree shrews are *lissencephalic* species similar to the mice, with a smooth cortex (*15*) and a GI value of 1.06. Tree shrews might represent an intermediate step in primate brain evolution. Therefore, we explored gene expression differences between fetal SVZ and other cortical layers in primate brains including the tree shrew. Our study revealed a previously unknown molecular regulator that plays a role in the gradual evolution in primate brains.

## Results

### CCNB1IP1 expression is human-specifically upregulated at SVZ

We used laser capture microdissection of the fetal cortices of tree shrews to generate transcriptome data from the cortical plate (CP), OSVZ, ISVZ, and VZ on embryonic Day 35 (E35), which is the peak stage of neurogenesis (**Fig.1C**). Principal component analysis (PCA) demonstrated that the CP (neuron-enriched layer) was completely separated from the GZ (progenitor-enriched layers). Within the GZ region, the VZ layers where APs resided differed from the ISVZ or OSVZ layers, which were enriched with BPs (**Fig.1D**). Consistent with the PCA results, the NPCs markers *SOX2*, *PAX6*, and *NES* and neuron markers *NEUROD6*(*39*), *TBR1*(*40*), and *FEZF2*(*41*) were highly expressed in the GZ and CP, respectively (**Fig.1D**). Therefore, the transcriptional signatures of the dissected layers reflect their spatial positions in the embryonic cortex of tree shrews. We further identified that tree shrew SVZ and expanded OSVZ are enriched for basal progenitors at E35 using SOX2 immunostaining (**Fig.1E**). Moreover, at E35, cortical plate is already consisted of upper layer SATB2^+^ neurons (*42*) and deeper layer TBR1^+^ neurons (**Fig.1E**).

Thus, we focused on the transcriptome evolution of the fetal SVZ, which is enriched in BPs and critical for neurogenesis and cortical evolution. We utilized fetal cortex lamina transcriptome datasets generated as previously described (*32, 43*) from representative species with different degrees of cortical folding, to identify genes whose expression was preferentially upregulated in the human SVZ. There were at least three biological replicates of each species (**fig. S1 A-D and Table S1**). In step 1, we identified the genes that were selectively expressed in SVZ regions compared with CP (**Fig.1F and fig.S1D; Table S2-3**). We then examined the differentially expressed genes (DEGs) between the ISVZ/OSVZ and CP and identified highly expressed genes for SVZ in each species. Finally, the expression of 153 genes was consistently significantly upregulated in the SVZ across all species (**Fig.1F and Table S3**). In step 2, we identified 1138 genes with specific expression in human SVZ. In step 3, we kept 541 genes with clearly orthologous in our comparative datasets. In steps 4 and 5, we evaluated these 541 orthologous genes with expression value FPKM (humans FPKM>50, other species FPKM<1.5). *CCNB1IP1* was the sole candidate gene that met these criteria (**Fig.1F and Table S4**).

### Human-specific upregulation of *CCNB1IP1* expression in neural progenitor cells

We investigated the patterns of *CCNB1IP1* expression during early brain development using data from previous studies (*3, 44, 45*). The findings about CCNB1IP1 highly expressed in human SVZ was corroborated by RNA-microarray data from human fetal cortex(*46*) especially at 15 weeks post-conception (15 wpc) (**fig.S1E**). Comparison results from macaque RNA-microarray expression data(*47*) showed that CCNB1IP1 expression was higher in CP (**fig.S1F**). Further, analysis with spatiotemporal transcriptome data from human fetal cortex (*48*) at 16wpc verified CCNB1IP1 expression pattern, although the difference between VZ and SVZ was indistinguishable (**fig.S1G**). Consistent with our findings in the fetal SVZ, *CCNB1IP1* was highly expressed in human neural progenitor RGCs, but was expressed at low levels in mouse RGCs (**fig.S1H**). Moreover, *CCNB1IP1* was highly expressed in human induced pluripotent stem cell (iPS)-derived NPCs compared to chimpanzee and bonobos NPCs (**fig.S1I**). Additionally, *CCNB1IP1* expression was remarkably upregulated in human brain organoids compared to gorilla brain organoids during organoid development (**fig.S1J**). These results solidified that *CCNB1IP1* expression increased in human NPCs during recent primate brain evolution.

We performed a re-analysis of single-cell RNA sequencing (scRNA-seq) data of the human fetal cortex (*31, 49*) to determine the cell types in which *CCNBI1P1* was expressed during early human brain development. The proportion of cells expressing this gene and the level of expression demonstrated that *CCNB1IP1* expression was particularly upregulated in NPCs (**fig.S2A-D**). According to the mitosis location, NPCs can be divided into APs and BPs(*50*). Therefore, we performed a comparative expression analysis of scRNA-seq data from micro-dissected VZ and SVZ regions at 16-18 wpc(*31*), a mid-fetal stage for human neurogenesis (**fig.S2E**). We used *PROM1* as a maker for APs(*31*) and *EOMES* for IPCs marker to identify APs cluster 3 (**fig.S2F**) Further, we used *MOXD1*, *HOPX,* and *LGALS3BP* as markers for BPs(*31*), and identified BPs cluster 1 (**fig.S2G**). Consistently, *CCNB1IP1* shows higher expression in BPs (cluster 1) than APs (cluster 3) (**fig.S2H**, *p* = 0012), indicating that *CCNB1IP1* is a BPs highly expressed gene. Moreover, at this stage, *CCNB1IP1* was already expressed in excitatory neurons not interneurons, suggesting *CCNB1IP1* mainly functions during neurogenesis for excitatory neurons (**fig.S2H**).

To further verify the human CCNB1IP1 expression pattern, we performed double immunostaining of cortices against SOX2 and CCNB1IP1 at 14 wpc of human. Similar to our findings above, most of the CCNB1IP1-expressed cells were co-localized with SOX2 at both VZ and SVZ (**Fig.2B**), and its expression is higher at SVZ according to the fluorescence intensity (**Fig.2B**, right panel). Additionally, at 14 wpc, CCNB1IP1 expression can be detected at CP (**Fig.2B**).

**Fig. 2.**
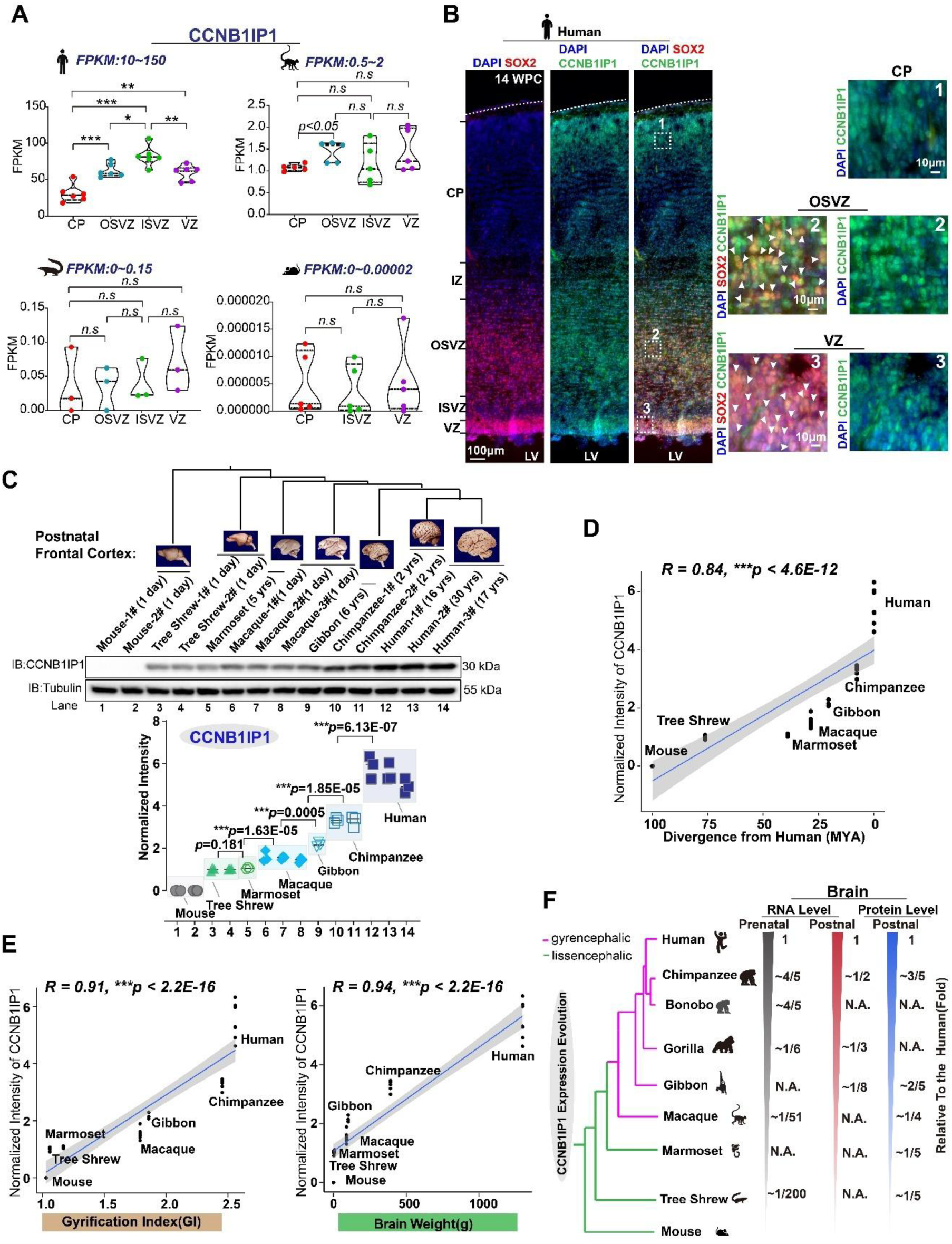
The expression of *CCNB1IP1* has gradually increased along primate brain evolution, coinciding with an increase in brain size and cortical folding. (**A**) Volcano plots demonstrating pairwise comparisons of gene expression between the laminae of humans, macaques, Chinese tree shrews, and mice. (**B**) (Left panel) Immunofluorescence staining of the cortices for SOX2 *and* CCNB1IP1 at 14 wpc of human. (Right panel) Area with white box shows high magnification of stained region including CP, OSVZ and VZ. (**C**) (top) Western blot analysis showing *CCNBIP1* expression patterns in human, chimpanzee, gibbon, macaque, marmoset, tree shrew, and mouse brains. (Bottom) Quantitative analysis of CCNBIP1 proteins. Each experiment was performed in triplicates (n=3 replicates for each sample, data represented as mean±SD, \**p* < 0.05, \*\**p* < 0.01, \*\*\**p* < 0.001, unpaired Student’s *t*-test). (**D**) Pearson’s correlation analysis showing a strong positive correlation between *CCNB1IP1* expression and evolutionary divergence. (**E**) Pearson’s correlation analyses demonstrating strong positive correlations between CCNB1IP1 expression and the cortical folding index (GI) and brain weight of primate species. (**F**) Schematic diagram showing the evolution of *CCNB1IP1* expression level.

Finally, to explore the human upregulation of *CCNB1IP1* in comparison to other species at the single-cell level, we performed an integration analysis using comparative scRNA datasets from human and chimpanzee organoids (*51*). This analysis demonstrated human-specific upregulation of *CCNB1IP1* expression, particularly in cycling NPCs at the G1/S phase (**fig.S3A-B**, cluster 5) and to a lesser extent, in non-progenitor cell types, including choroid and inhibitory neurons (**fig. S3A-C**). A similar integration analysis using human and macaque fetal cortices revealed broad upregulation of *CCNB1IP1* expression in humans **(fig.S3D-F)**. Analysis of another organoid scRNA-seq(*52*) confirmed that *CCNB1IP1* expression was highly upregulated in human NPCs during organoid development compared with chimpanzees and macaques (**fig.S3G-H**).

### Gradual increase of neuronal *CCNB1P1* expression during primate brain evolution

Utilizing spatial and temporal human brain transcriptome data (*53*), we found that human *CCNB1IP1* was highly expressed at the prenatal stage, and its expression gradually decreased after birth, showing comparable levels in different brain regions (**fig.S4A-B**). We also observed recent human-specific upregulation of *CCNB1IP1* in a study of eight adult brain regions (*54*) (**fig.S4C**). *CCNB1IP1* expression was detected in diverse cell types in adult human brains, especially in neurons. Contrastingly, CCNB1IP1 was lowly expressed in adult chimpanzee and macaque brains (**fig.S4D-E**). These results indicate that *CCNB1IP1* expression has recently increased in both NPCs and neurons during human brain evolution.

To corroborate this conclusion and elucidate the evolutionary changes in CCNB1IP1 at the protein level, we performed western blotting (WB) analyses of the cortices of seven representative primate species and non-primate outgroups (**Fig.2C** top panel). This analysis confirmed that CCNB1IP1 expression gradually increased toward its highest expression in the human brain in primates (**Fig.2C** bottom panel) and that CCNB1IP1 expression was strongly correlated with divergence times (**Fig.2D**, *R* = 0.84, *p* < 4.6E-12). Further, CCNB1IP1 expression was significantly and positively correlated with brain weight and degree of cortical folding (**Fig.2E**, *R* = 0.91, and *R*=0.94, respectively, *p<*2.2E-16 for both comparisons). Notably, the small New World monkey, the marmoset, which is a lissencephalic species with a smooth cortex, showed comparable CCNB1IP1 expression levels with tree shrew. However, these were significantly lower than macaque, a gyrencephalic species with a folding cortex (**Fig.2C**). We summarized the expression levels of human and nonhuman primate brains (*3, 32, 43–45, 54*) and observed that CCNB1IP1 expression gradually increased during human brain evolution (**Fig.2F**).

### Cis regulatory evolution of CCNB1IP1 expression

The spatial and temporal expression of a gene is regulated by *cis-* and *trans*-genetic factors, as well as other non-genetic factors (*55*). Here, we examined scRNA-seq data from human-chimpanzee hybrid organoids (*56*). The expression divergence of *CCNB1IP1* between humans and chimpanzees was consistent with the effects of large *cis-* factors (**Fig.3A**). Thus, we examined cis regulatory elements (CREs) located in the upstream region of *CCNB1IP1* gene using 3D epigenome data of human fetal cortex (*57*). We found two potential CREs located in the promoter region of human *CCNB1IP1* gene carrying the peaks for open chromatin states based on the Assay for Transposase-Accessible Chromatin with high-throughput sequencing (ATAC-seq) data in the main cell types of human fetal brain (**Fig.3B**). To further strengthen this result, we examined published chromatin modification data including H3K4me3-promoter histone marker, H3K27ac-enhancer histone marker et al during the human NPC differentiation process(*58*). The results showed that the identified CRE1 but not CRE2 carried the peak of H3K4me3 in both human NPCs and neurons (**fig.S5A-B**), the same location with ATAC peak as shown in Fig. 3B, indicating that CRE1 is capable of regulating activity as promoter. As expected, compared with differentiated neurons, the H3K4me3 peak height of CRE1 was much higher in human NPCs (**fig.S5B**), which is consistent with high expression of CCNB1IP1 in NPCs. However, for CRE2, we did not observe histone modification peak in either NPCs or neurons (**fig.S5A-B**).

**Fig. 3.**
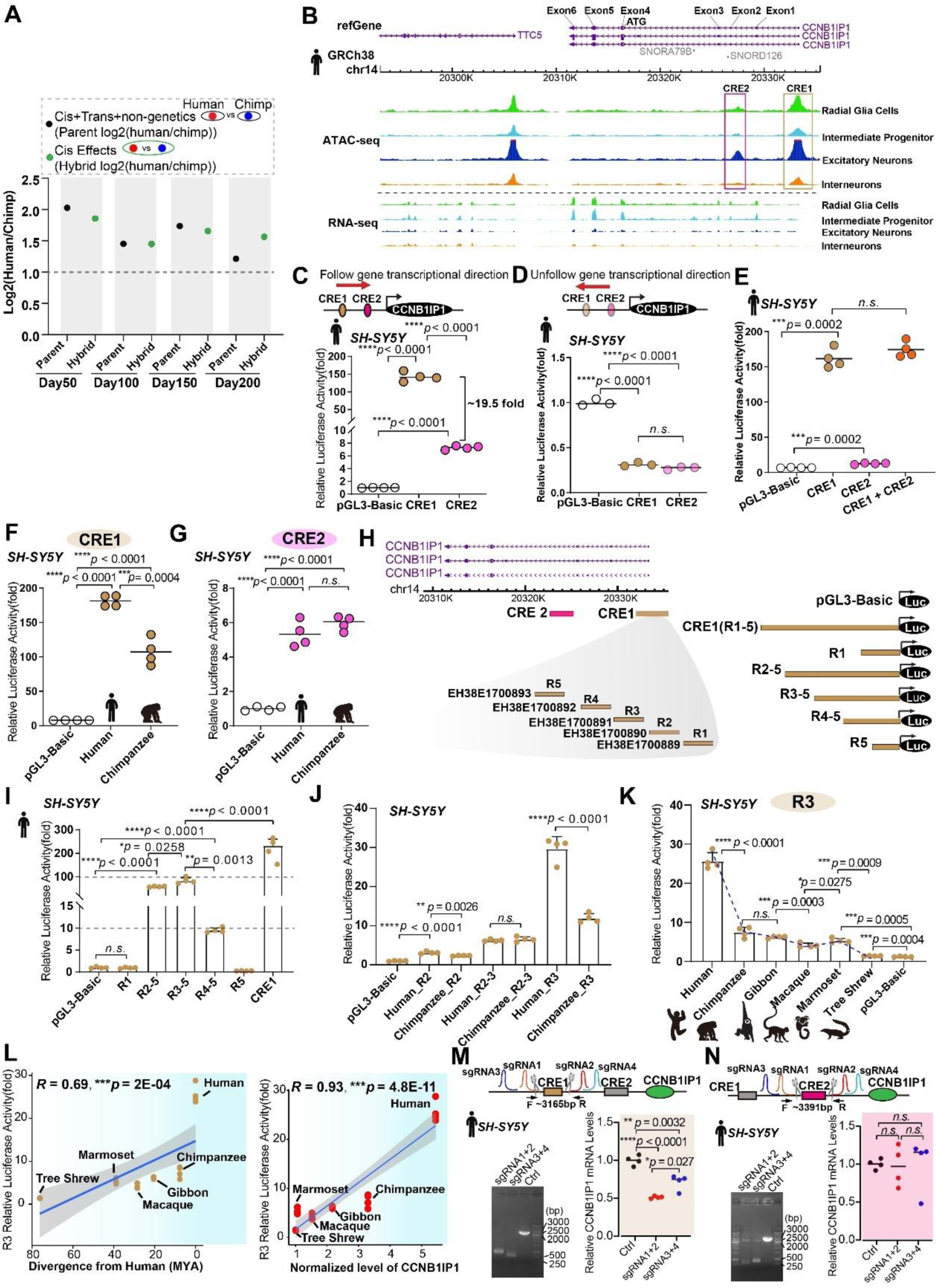
Disentangling *cis* effects for regulating human *CCNB1IP1* expression. (**A**) Dot plot showing cis and trans divergence regulating *CCNB1IP1* expression between parental lines and hybrids during cortical spheroid (CS) development. Parents: Three human and three chimpanzee iPS lines inducing CS differentiation. Hybrid: five human–chimpanzee hybridized iPS cell lines inducing CS differentiation. Log2(human/chimp), with positive values indicating higher expression in humans than in chimpanzees. \**CCNB1IP1* expression data were downloaded from Agolia et al(*56*). (**B**) Cis-regulatory element CRE1 and CRE2 are at the promoter region of *CCNB1IP1* gene with ATAC signal peak. (**C**) (Top) Both CRE1 and CRE2 are cloned into the luciferase reporter following the CCNB1IP1 transcription direction and (bottom) subsequent reporter analysis of their activation activities. (**D**) (Top) Both CRE1 and CRE2 are cloned into the luciferase reporter in the opposite direction of CCNB1IP1 transcription and (bottom) subsequent reporter analysis of their activation activities. (**E**) Luciferase reporter analysis testing synergistic effects of CRE1 and CRE2 grouped together. (**F**) Luciferase reporter analysis of human and chimpanzee CRE1 activation activities. (**G**) Luciferase reporter analysis of human and chimpanzee CRE2 activation activities. (**H**) (Left) A schematic diagram showing CRE1 contains five annotated regulatory elements. (Right) A schematic map of the 5’ deletion constructs of CRE1. (**I**) Luciferase reporter analysis of the 5’ deletion construct activities. (**J**) Luciferase reporter analysis of R2, R3 and R2-R3 activities between human and chimpanzee species. (**K**) Luciferase reporter analysis of R3 activities among human, chimpanzee, gibbon, macaque, marmoset and Chinese tree shrew species. The trend line is indicated with blue color. (**L**) (Left) Pearson’s correlation analysis demonstrating a strong positive correlation between R3 activation activity and evolutionary divergence. (Right) Pearson’s correlation analysis demonstrating a strong positive correlation between R3 activation activity and CCNB1IP1 expression level. (**M**) (Left) The results of running gel analysis on selected clones after CRISPR-Cas9 cutting with designed paired sgRNAs for CRE1, as indicated. (Right) Quantification of CCNB1IP1 RNA expression levels following CRE1 deletion. (**N**) (Left) The results of running gel analysis on selected clones after CRISPR-Cas9 Cut with designed paired sgRNAs for CRE2 as indicated. (Right) Quantification of CCNB1IP1 RNA expression levels following CRE2 deletion. All statistical data are presented as mean or mean ±SD, \**p* < 0.05, \*\**p* < 0.001, \*\*\**p* < 0.001, *n.s*, not significant, as determined using the unpaired Student’s *t*-test.

Interestingly, we found that neither of these two CREs was intact in the mouse genome comparing with primates (**Fig.S5C**), which is consistent with the low expression of CCNB1IP1 in the mouse brain (**Fig.2C**). However, both CREs can be detected in other mammalian species as early as in platypus, an extant species of egg-laying mammals, implying a complicated evolutionary history of CREs for CCNB1IP1.

CREs are located at 5’ upstream of CCNB1IP1 and can be functional as promoters or enhancers. To explore it, we cloned human CRE1 and CRE2 into a pGL3-basic vector following the direction of gene transcription and performed luciferase reporter assays. Both CRE1 and CRE2 showed significantly upregulated activities compared with control suggesting that the identified CREs were functionally regulatory elements (**Fig. 3C**). However, when CRE1 and CRE2 are cloned into pGL3 in reserve direction unfollowing gene transcription direction, the regulatory activities are dramatically decreased (**Fig.3D**), indicating that both CRE1 and CRE2 were promoters. Moreover, consistent with the higher peak of ATAC and H3K4me3 (**Fig.3B and fig.S5A-B**), the CRE1 activity was much higher than CRE2 by ∼19 fold (**Fig.3C**). Then, we wondered whether there is a synergistic effect by grouping them together. However, the report assay results showed that adding CRE2 did not show enhanced effects compared with CRE1 alone (**Fig.3E**), which indicates that CRE1 is the main regulatory element for CCNB1IP1.

We then compared the regulatory activities of human and chimpanzee CREs, and found that human CRE1 activity was significantly higher than that of the chimpanzees (**Fig.3F**). The activities of CRE2 were comparable between human and chimpanzee (**Fig.3G**). In fact, according to the human ENCODE data (https://www.encodeproject.org/), human CRE1 can be further annotated into five regulatory elements (**Fig.3H**, R1-R5). To determine which regulatory element of CRE1 is responsible for the activity, a series of fragments with different R1-R5 deletions were generated by PCR and ligated to the pGL3 basic vector (**Fig.3H**, right panel). We found that the activity of CRE1 was mainly driven by R2 and R3, as constructs harboring those two elements exhibited the highest activities (**Fig.3I**). Focusing on R2 and R3 and by comparing activities with chimpanzee elements, we found that human R3 exhibited the higher and more divergent regulatory activities compared to R2 (**Fig.3J**). We then examined the regulatory activities of R3 from four representative species including gibbon, macaque, marmoset and tree shrew, by cloning their R3 elements into pGL3 basic vector. We observed that 1) the human R3 exhibited highest activity among the tested species (**Fig.3K**). 2) the activity of R3 was significantly upregulated in all the tested species and exhibits a gradually reducing trend from human to tree shrew (**Fig.3K**, blue broken line). Further, the regulation activity of R3 was significantly and positively correlated with both divergent times (**Fig.3L** left panel, *R* = 0.69, *p=*2E-04) and CCNB1I1P expression level (**Fig.3L** right panel, *R* = 0.93, *p=*4.8E-11), indicating that the evolution of R3 regulatory activity is responsible for the increased expression of *CCNB1IP1*.

Finally, to examine the regulatory effects of CREs on CCNB1IP1 expression, we performed CRISPR-Cas9 experiments using paired sgRNAs to cut CRE1 and CRE2, respectively, in *SH-SY5Y* cells. Consist with the results of regulatory activities, we found that following CRE1 deletion, the expression of *CCNB1IP1* was significantly downregulated (**Fig.3M**, 48.5% downregulated with sgRNA1&2; 28.6% downregulated with sgRNA3&4), suggesting that CRE1 is capable of regulatory function. However, for CRE2, consistent with its low regulatory activity (**Fig.3C**), the expression of CCNB1IP1 shows no difference with controls (**Fig.3N**) following CRE2 deletion, again suggesting that CRE2 has limited capacity for regulating function.

Together, our results show that the evolution of CRE1 is associated with the changes in the expression of *CCNB1IP1* during the origin and evolution of primates.

### CCNB1IP1 promotes proliferation of BPs through shortening the length of G1 phase

Based on the results of CCNB1IP1 expression and related CREs evolution, we wonder whether the CCNB1IP1 gene is also under rapid evolution. The protein sequence alignment across 20 mammalian species indicated that CCNB1IP1 is relatively conserved from antbear to humans (**fig.S6A**). Further, pairwise comparisons showed that the similar percentage was as high as above 0.88 (**fig.S6B**), confirming that CCNB1IP1 is a highly conserved protein. However, it was expressed at extremely low levels in the brains of mice, contrary to its abundant expression in the brains of primates (**Fig.2C)**. This indicates that *CCNB1IP1* may have evolved a new role in the primate brain.

CCNB1IP1 is mainly expressed in progenitor cells and the primary function is to regulate meiotic recombination in mammals (*59–61*), implying that CCNB1IP1 may regulate cell cycle during primate brain evolution. Firstly, we verified that CCNB1IP1 overexpression significantly increased the proportion of proliferating cells (**fig.S7A,** 12.29% vs. 18.18%, *two-tail unpaired t-test p* = 0.00012). Then, to investigate whether *CCNB1IP1* could influence the cell cycle status, we performed flow cytometry using propidium iodide, a DNA-labelling dye, to quantify the cell cycle phase of N2A cells with and without CCNB1IP1 transfection. The results suggested that CCNB1IP1 overexpression modestly yet significantly reduced the number of cells in the G1 phase and increased that in the S phase (**Fig.4A**). Furthermore, we performed fluorescent ubiquitination-based cell cycle indicator (FUCCI)(*62*) imaging to trace the G1 phase at 24, 36, and 48 h. The proportion of G1 phase cells and the cell doubling time, which is an indicator of G1 length, were significantly reduced following CCNB1IP1 transfection (**Fig.4B**). To examine whether CCNB1I1P overexpression could promote NPCs proliferation, we performed CCNB1IP1-in utero electroporation (IUE) *in vivo* at E13.5, injected EdU at E14.5 for 24 h, and analysed it at E15.5 (**Fig.4C**). CCNB1IP1 specifically promoted the proliferation of progenitor cells in the SVZ (**Fig.4D**, bottom panel, EdU^+^GFP^+^/GFP^+^ from 17.91% to 34.78%, *two-tail unpaired t-test p* = 0.0049). Consistently, the number of cycling cells (Ki67^+^) was increased in the SVZ of CCNB1IP1 electroporated regions at E16.5 by performing IUE at E14.5 (**Fig.4E**).

**Fig. 4.**
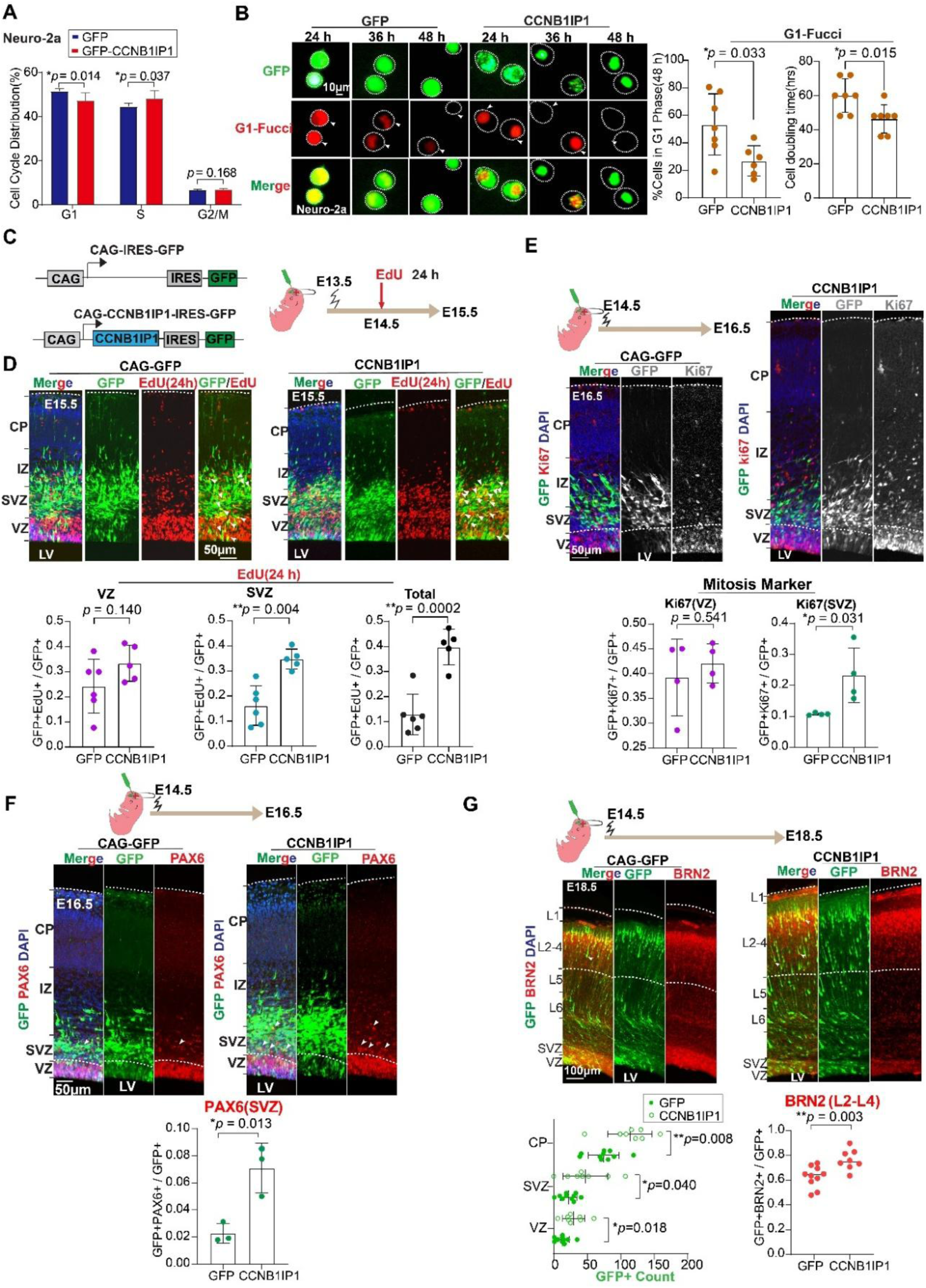
The expression of *CCNB1IP1* promotes BPs proliferation, and neurons output in mice. (**A**) Cell cycle distribution analysis after 48 h transfection of GFP and CCNB1IP1-GFP in neuro-2a cells. (n=5-6 replicates, data represented as mean±SD, \**p* < 0.05, unpaired Student’s *t*-test). (**B**) Left panel: Representative images showing separate transfection of GFP and *CCNB1IP1* (green) with G1-Fucci (red) constructs at 24, 36, 38 h. Right panel: Quantification of GFP^+^ and CCNB1IP1-GFP^+^ co-localization with G1-Fucci^+^. (n=6-7 replicates, data represented as mean± SD, \*\**p* < 0.01, unpaired Student’s *t*-test). (**C**) Top panel: Electroporation of CAG-GFP and CAG-CCNB1IP1-GFP constructs at E13.5, followed by EdU at 24 h and analysis at E15.5. *White arrowheads indicate GFP^+^Edu^+^ double-positive cells. (**D**) Quantification of GFP^+^ and CCNB1IP1-GFP^+^ colocalization with EdU^+^ in the VZ, SVZ, or VZ + SVZ. (n=5-6 embryos from two IUE experiments, data represented as mean±SD, \*\**p* < 0.01, unpaired Student’s *t*-test). (**E**) Representative images of the mitosis marker Ki67 and quantification of Ki67 ^+^ in GFP^+^ cells, (n=4 embryos from one IUE experiment, data represented as mean± SD, \**p* < 0.05, unpaired Student’s *t*-test). (**F**) Representative images of the NPCs marker PAX6 immunofluorescence after electroporation with GFP or CCNB1IP1 at E14.5. Quantification of PAX6^+^ in GFP^+^ cells in the VZ and SVZ (n=3 embryos from one IUE experiments, data represented as mean± SD, \**p* < 0.05, unpaired Student’s *t*-test). (**G**)(Top) Representative images of the upper layer neuron marker BRN2 at E18.5, following electroporation of GFP or CCNB1IP1 at E14.5. (Bottom) Quantification showing the effects of *CCNB1IP1* expression on the number of GFP^+^ and BRN2^+^ cells. Arrows indicate GFP^+^BRN2^+^ cells (n=9 embryos from three IUE experiments, data represented as mean± SD, \**p* < 0.05, unpaired Student’s *t*-test).

More than that, forced expression of CCNB1IP1 significantly increased the proportions of the BP marker-Pax6^+^(SVZ) by 3.12-fold (**Fig.4F**, *two-tail unpaired t-test p* = 0.013). To validate these findings, we performed additional IUEs at E13.5 in mouse embryonic neocortices and analyzed the electroporated region (left side, L) at E15.5, using a non-electroporated region (right side, R) as a control (**fig.S7B**). CCNB1IP1 significantly increased the number of BPs in the electroporated region compared with that in the control regions (**fig.S7C-E**). These results show that CCNB1IP1 promotes BP (including bRGC and IPCs) proliferation in the SVZ. Because the number of BPs is associated with the output of neurons during neurogenesis (*19*), we performed long-term IUE of CCNB1IP1 at E14.5, followed by analysis at E18.5. Quantification of Brn2 in upper layer (L2-L4) neurons showed that CCNB1IP1 significantly increased Brn2^+^GFP^+^ neurons by 13.7% (**Fig.4G**, *two-tail unpaired t-test p* = 0.0033). Therefore, CCNB1IP1 overexpression can increase the neuron output during neurogenesis.

### CCNB1IP1 knock-in mice developed folding cortices and cognitive enhancement

To determine the functional effects of CCNB1IP1 *in vivo*, we generated CCNB1IP1 conditional knock-in mice (cKI) driven by the CAG promoter (**fig.S8A-B**). To conditionally express CCNB1IP1 in NPCs, we used *Emx1-Cre* mice, a transgenic line that mediates recombination events in cortical NPCs, starting at E10.5(*63*). Western blotting confirmed elevated levels of CCNB1IP1 in the brains of cKI*^Emx1-cre^* mice, whereas the controls showed no CCNB1IP1 expression (**fig.S8C**).

Analysis of the cortical phenotype of cKI*^Emx1-cre^* on postnatal Day 1 (P1) revealed significant folding of the neocortex in *Emx1-cre*-cKI (**fig.S8D**). Local GI was significantly elevated in cKI*^Emx1-cre^* mice compared with the control (1.11 vs. 1.00, *p* = 0.00093, **fig.S8D**). We further examined the layer structure of cKI*^Emx1-cre^*to assess the contribution of layer neurons to cortical folding. Although layer formation was retained, CCNB1IP1 expression significantly increased the number of Brn2^+^ neurons but not Tbr1^+^ neurons (**fig.S8E**), indicating that the upper layer neurons contributed to cortical folding.

We performed additional knock-in experiments by generating *Nestin-cre-*cKI mice, starting recombination events in the neural stem cells at E11.5(*64*). We verified that CCNB1IP1 was highly expressed in the cortices of cKI*^Nestin-cre^* mice (**Fig.5A**). Furthermore, we examined the phenotypes of cKI*^Nestin-cre^* mice, and the brain size of cKI*^Nestin-cre^* mice was 11.02% larger than that of controls (**Fig.5B**, *two-tail unpaired t-test p* = 0.022). Similarly, CCNB1IP1 expression significantly increased the number of Brn2^+^ neurons (**Fig.5C**). Cortical folding developed in cKI*^Nestin-cre^* mice (**Fig.5D-E**), resulting in significant changes in local GI (**Fig.5F**, 1.07 vs. 1.00, *p* < 0.0001). To further explore whether the cellular mechanisms were the same as those of IUE, we analyzed the E15.5 embryonic cortex of cKI*^Nestin-cre^* mice. The embryonic cortex size was significantly expanded (**fig.S8F**). We also confirmed an increase in BPs in the SVZ of cKI*^Nestin-cre^* mice (**fig.S8G-I**). These findings support the hypothesis that CCNB1IP1 is sufficient to increase BP proliferation.

**Fig. 5.**
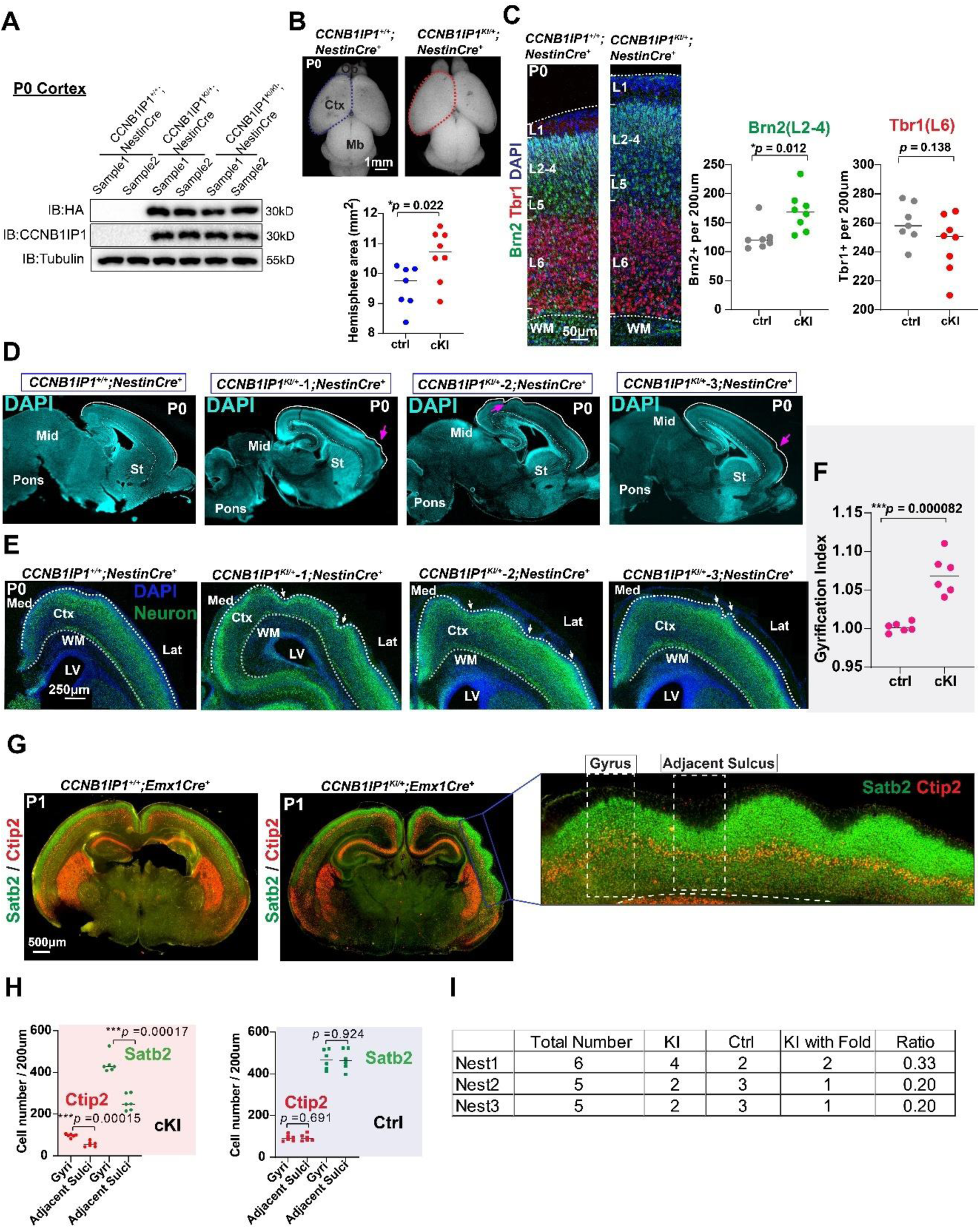
*CCNB1IP1* conditional knock-in mice exhibit cortical expansion and folding pattern. (**A**) Western blotting analysis showing CCNB1IP1 upregulation in the P0 cortex of CCNB1IP1Nestin-Cre KI mice. (**B**) Dorsal view of WT and CCNB1IP1Nestin-Cre cKI P0 brains from the 2 nests. (ctrl n=7 mice, cKI n=8 mice, data represented as mean, \**p* < 0.05, unpaired Student’s *t*-test). (**C**)Cortical layer marker analysis revealed a significant increase in BRN2+(L2-4) but no significant changes in TBR1+(L6) in cKI. (ctrl n=7 mice, cKI n=8 mice, data represented as mean, \**p* < 0.05, unpaired Student’s *t*-test). (**D**)Representative images of cortical folding (pink arrows) in cKI Nestin-cre mice at P0. (**E)**Neuronal immunostaining showing a cortical folding pattern (arrowheads) in Cki. (**F**)and comparison of local gyrification indices in cKI. (ctrl n=6 mice, cKI n=6 mice, data represented as mean, \*\*\**p* < 0.001, unpaired Student’s *t*-test). (**G**)Representative images of the cortical layer marker SATB2 (L2-4) and CTIP2 (L5) in cKI mice at P0. Area with blue rectangle to be magnified to show gyri and adjacent sulci structure with white rectangle in folding locations. (**H**)Quantification of STAB2 and CTIP2 between the gyri and adjacent sulci structure in both cKI and control mice. (ctrl n=6 mice, cKI n=6 mice, data represented as mean, \**p* < 0.05, unpaired Student’s *t*-test). (**I**)The percentages of cortical folding from 3 independent nests at P1 age.

Further, we examined the layer neuron distribution by immunostaining with Satb2 as a marker for the upper layer and Ctip2 for the deep layer (**Fig.5G and fig. S9**). Cortical layers follow the cortical folding trend and are well preserved between the gyrus and adjacent sulcus structure (**Fig.5G and fig.S9)**, which is consistent with the true folding phenotype that occurs only in all cortical layers not entire cortical walls(*65*). Further, layer neurons are significantly reduced in sulci, especially for upper layer Satb2 neurons, compared with gyri (**Fig.5H**), which is also similar with the layer distribution in gyrencephalic species e.g. macaque (*66*). In fact, not every cKI mice carry cortical folding phenotype and about 20%-33% of cortical folding per nest computed from three independent nests (**Fig.5I**).

Finally, we conducted sequential behavior experiments according to the degree of external stimulus pressure to investigate whether CCNB1IP1-KI mice with folding cortex exhibited distinctive cognitive behavior compared with controls. In the open field test (*67*), cKI*^Emx1-cre^*mice tended to travel longer distances from the center and at faster speeds. However, they spent significantly less time staying or resting, which is interpreted as reflecting an increased level of curiosity and anxiety (**Fig.6A**). In the novel object recognition test (*68*), cKI*^Emx1-cre^* mice were prone to discriminate between familiar and novel objects and spent significantly longer time with the novel objects than control mice, implying that recognition and memory abilities are possibly increased (**Fig.6B**). In the three-chamber social test (*69*), cKI*^Emx1-cre^* mice did not show any difference from the controls (**Fig.6C**). However, in the Morris water maze test (*70*), cKI*^Emx1-cre^* mice spent less time and travelled shorter distances to reach the platform than the control group following training (**Fig.6D**). This trend was evident on day 3, the middle day of training, and the cKI mice exhibiting better learning abilities than control mice (**Fig.6D**). These results suggest that changes in the brain size and the degree of cortical folding, especially for species carrying a smooth cortex, may have an impact on brain function, including learning and memory.

**Fig. 6.**
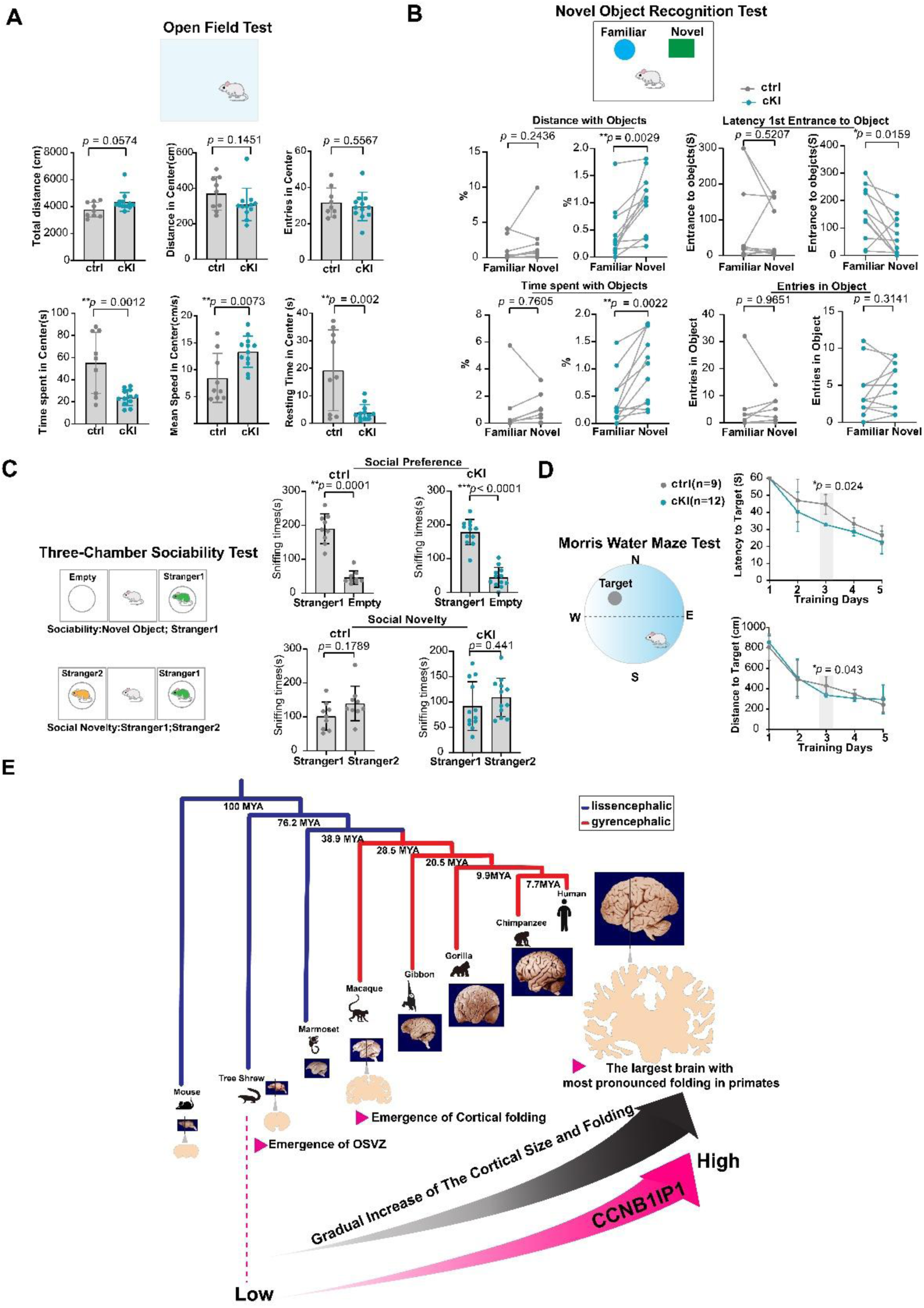
*CCNB1IP1* conditional knock-in mice exhibit enhanced cognitive ability. (**A**) Open-field assay showing cKI curiosity and anxiety (control, n = 9; cKI, n = 12). (**B**) Novel object recognition test results, demonstrating cognitive abilities in cKI mice (control, n = 9; cKI, n = 12). (**C**) Three-chamber sociability test results demonstrating social behavior between controls and cKI mice (control, n = 9; cKI, n = 12). (**D**) Morris water maze test results showing enhanced learning but not memory abilities in cKI (control, n = 9; cKI, n = 12). (**E**) The proposed model for CCNB1IP1 regulation of cortical size and folding during primate evolution. During primate brain evolution, cortical size and folding have gradually increased, reaching the peak development observed in the human species (shown as a gray gradient). As a consequence, the expression of *CCNB1IP1* gene gradually increased within the cortex, commencing before the origin of primates and notably intensifying from the tree shrew onwards, reaching its highest level in the human species (shown as a ruby gradient). This evolutionary interaction coincides with an increase in the proliferation abilities of neural progenitors in primate brains, ultimately contributing to the formation of folded cortex during primate brain evolution. Notably, among primates, the human brains display the highest expression of CCNB1IP1 and the most expanded and folded cortex. *The ruby-colored arrowhead emphasizes key events in the timeline of the primate brain evolution. All statistical data are presented as mean or mean ±SD, \**p* < 0.05, \*\**p* < 0.001, \*\*\**p* < 0.001, \*\*\*\**p* < 0.001, n.s, not significant, as determined using an unpaired Student’s t-test.

## Discussion

Previous molecular studies of human brain evolution have revealed the contributions of human-specific genes generated by DNA duplication (*2–7*), those originated *de novo* (*8*), great ape-specific gene *TBC1D3* (*28*) and primate-specific gene *TMEM14B* (*29*). However, to our knowledge, none of these studies can explain the gradual increase of cortical size and folding along primate brain evolution. As many features of the human brain have long evolutionary origins, it is important to understand the contribution of conserved genes to human brain-specific phenotypes. In this study, through cross-species comparisons, we identified CCNB1IP1 as a key regulator of cortical evolution. CCNB1IP1 expression gradually increased with primate brain evolution, which is implicated in the shortening of the G1 phase and an increase in the proliferative abilities of BPs, indicating that high levels of CCNB1IP1 may promote G1 /S transition process. These processes contribute to an increase in neuronal output and ultimately promote cortical size and folding increase (**Fig.6E**).

We demonstrated that the evolution of *CCNB1IP1* expression is regulated by the upstream *cis*-regulatory elements, indicating that the evolution of CRE might be linked to the evolution of *CCNB1IP1* expression. However, only CRE1 is capable of CCNB1IP1 transcriptional activation, implying that CRE2 regulatory function may be not as promoters but as other regulatory elements, which warrant further study in future. Moreover, the activity of human CRE1 was significantly higher than that of the chimpanzee CRE1, indicating an additional enhancement of cis-regulatory element in the recent evolutionary history of human brain. Our serial deletion experiments further suggested that the evolution of cis-regulatory activity of R3 element within the CRE1 mirrors the gradual increase of CCNB1IP1 and the corresponding evolution of brain size and cortical folding. Future functional studies are required to elucidate the specific R3 mediated trans-regulatory mechanism involved in the regulation of *CCNBIIPI*.

Surprisingly, CCNB1IP1 expression has also been detected in the brains of tree shrews, which are a close outgroup of primates with intermediate brain phenotypes, including implications for the emergence of OSVZ (*33–35*). Focusing on the key germinal region SVZ for cortical expansion and folding in primates (*18*) using Chinese tree shrews, and through multiple species comparisons, enabled the systematic identification of human-specific upregulated genes in SVZ, such as *CCNB1IP1*. Consistently, the induction of cortical folding in the mouse brain suggests that CCNB1IP1 may have played an unrecognized role in the regulation of cortical folding during primate evolution. Finally, behavioural tests revealed improved memory and cognitive abilities in KI mice compared with the control, indicating that CCNB1IP1-induced cortical expansion and folding may affect cognition.

This study revealed that *CCNB1IP1* expression accelerates G1/S phase transition and the subsequent proliferation of BPs. Shortening of the G1 phase was previously shown to increase the rates of proliferative divisions at the expense of differentiation, which results in the expansion of OSVZ in primates (*24–26*). A recent genome-wide CRISPR interference screen in human and chimpanzee stem cells revealed that human-specific changes were enriched in gene networks that promote G1/S phase progression (*71*). These observations indicate that cell cycle regulation may have played an important role in human brain evolution.

In summary, cortical expansion and folding are complex phenotypes involving the sophisticated coordination of molecular, cellular, and mechanical processes during brain development (*19*). Our work identified cis-regulatory evolution of CCNB1IP1 that governs the gradual evolution of the neocortex in primates, thus advancing our understanding on the origin of human intelligence.

## Materials and Methods

### Ethics Statement

Animal research using Chinese tree shrews and mice was conducted according to the guidelines approved by the Animal Care and Use Committee of the Kunming Institute of Zoology, Chinese Academy of Sciences. The animals involved in this study were all from standard feeding environments and did not carry potential pathogens or parasites that would cause great interference to the results of the scientific experiments. Chinese tree shrews were obtained from the Laboratory Animal Center of the Kunming Institute of Zoology. C57BL/6J mouse Cre lines *Nestin*-Cre(*64*) and *Emx1*-Cre(*63*) were from Jackson Laboratory (Bar Harbor, ME, USA).

### Tissue Samples

The frozen cerebral cortexes of three humans were obtained from the Chinese Brain Bank Center (Wuhan, China). The donors of all tissues had no known neuronal diseases or history of drug abuse. Two chimpanzees, one gibbon, three rhesus macaques, one marmoset, two Chinese tree shrews, and two mice were used in this study. These tissue samples were obtained from the Kunming Institute of Zoology. They were healthy and free of pathogens. The experiments were approved by the internal review board of the Kunming Institute of Zoology.

Human embryo cortical samples at post-conception weeks (PCW) 11, 14, and 15 were acquired from Baoding Second Central Hospital and used for research with the donor’s consent. The fetal cortex was freshly microdissected and flash-frozen in liquid nitrogen for subsequent experiments. All experimental procedures were reviewed and approved by the Medical and Institutional Ethics Committee (IGDB-2020-IRB-001).

### Chinese tree shrew embryo acquisition

To obtain embryos on the exact days of development, female and male tree shrews were kept in the same cage overnight and separated from each other the next day. The day of separation was considered embryonic day 0 (E0). After 10 days, trained veterinarians gently touched the uterine position of the female tree shrews to determine whether they were pregnant. The exact number of days of development was determined following these procedures.

### Conditional knock-in mouse models

A CCNB1IP1 conditional knock-in mouse model with flox sites was generated using CRISPR/Cas9. The gRNA to mouse ROSA26 gene, a donor vector containing "CAG promoter-loxP-PGK-Neo-6*SV40pA-loxP-Kozak-Human CCNB1IP1 CDS-HA tag-rBG pA" cassette, and Cas9 mRNA were co-injected into fertilized mouse eggs to generate targeted conditional knock-in offspring. The founder (F0) animals were confirmed using PCR, followed by sequence analysis. They were then bred into wild-type mice to test germline transmission and generate F1 animals. The HA tag is located upstream of the TGA stop codon of human *CCNB1IP1*. Five pups were identified as F0 using PCR genotyping of genomic DNA collected from the tails. F0 mice were continuously bred with wild-type C57BL/6 mice. The correct gene targets in F1 animals were confirmed through Southern blotting of tail DNA samples. CCNB1IP1-cKI mice were generated by crossing CCNB1IP1^flox/+^ mice with nestin-Cre or Emx1-Cre mice. CCNB1IP1-cKI mice (CCNB1IP1^flox/+^; Nestin-Cre) were identified through genotyping. The oligonucleotide primers used for genotyping are listed in Table S5.

### Cell lines

HEK293 cells were cultured in Dulbecco’s Modified Eagle’s medium (DMEM) (Gibco, C11995500BT) supplemented with 10% fetal bovine serum (FBS) (Gibco, A5669401), penicillin, and streptomycin (100 U/mL) (Gibco, 15140122). SH-SY5Y cells were cultured in DMEM containing 10% FBS, 2% GlutaMAX, and 1% penicillin-streptomycin at 37°C with 5% CO_2_. Neuro2A cells were cultured in DMEM containing 10% FBS, 1% glutamine (Gibco, 25030081), 1% NEAA (Gibco, 11140050), and 1% penicillin-streptomycin. The cells were transfected using Lipofectamine 3000 (Invitrogen, L3000008) according to the manufacturer’s instructions.

### Laser Capture Microdissection (LCM)

Fresh tissues were slowly frozen in an OCT compound (Sakura, 4583) with a mixture of dry ice and alcohol. Sections were cut at 20–25 µm on a cryotome and attached to membrane slides (Leica, 11505189) with a 2 µm PEN-Membrane. DEPC-treated water was used in all solutions to prevent RNA degradation. To obtain a 1% Nissl staining solution, 1 g crystal violet (Sigma-Aldrich, C5042) was dissolved in 100 mL DEPC-treated water (Biosharp, BL510A) overnight on a shaker at room temperature about 25-35℃. Subsequently, filter paper was used to remove insoluble impurities. Tissue sections were dehydrated with 50%, 75%, and 100% gradient alcohol for 1 min. Dehydrated sections were stained with 1% cresyl violet acetate solution until the cell color turned purple-blue. The target tissue area was cut using a laser microdissection system (Leica, LMD7000) and collected into 0.6 mL microtubes (Axygen, MCT-060-C) containing TRIzol (Invitrogen, 15596026). RNA was isolated using the miRNeasy Micro Kit (QIAGEN, 217084).

### RNA Sequencing Library Construction

RNA sequencing libraries were constructed using a TruePrep RNA library prep kit for Illumina (TR502-02, Vazyme). Briefly, mRNA was purified using capture beads with Oligo-dT and eluted from the beads for fragmentation. The first cDNA strand was synthesized using random primers, whereas the second strand was synthesized through digestion with RNase H. Clean DNA beads (Vazyme, N411) were used for size selection following end repair and adapter ligation. The length of the final RNA sequencing library was 320–420bp. An Agilent high sensitivity DNA Kit (Santa Clara, 5067-4626) was used for quality tests.

### Luciferase reporter assays

All transfections were performed in 24-well plates (NEST, 702001) with at least three replicates. Approximately 2 × 10^5^ cells were seeded 24 h before transfection. Equal numbers of cells were plated in the 24 and 6-well plates and grown to 80% confluence. Next, vectors were mixed in Opti-MEM (Gibco, 11058021) with lipofectamine 3000 (Invitrogen). The solution was incubated for 30 min at room temperature and added to cultured cells. After 4–6 h, the medium was changed to DMEM (Gibco) containing 10% FBS. Cells were grown in 24-well plates and transfected with the vectors, with pTK-*Renilla* as an internal control, using Lipofectamine 3000. Luciferase activity was measured 28–32 h after transfection. Luciferase activity in the cell extracts was determined using a Dual-Luciferase Reporter Assay System (Promega, E1910) according to the manufacturer’s instructions. Relative light units were measured using a luminometer. Each experiment was repeated at least three times.

### Cloning of CCNB1IP1regulatory regions among different species

Human and chimpanzee fragments of the CRE1 ∼1,226bp and CRE2 ∼2001bp were obtained by PCR amplifications. The amplicons were cloned into the pGL3 basic firefly luciferase vector (Promega). *XhoⅠ* and *HindⅢ* restriction sites were introduced in the forward and reverse primers, respectively and employed for cloning into the luciferase reporter gene plasmid. The amplified DNA fragment was digested with *XhoⅠ*/*HindⅢ* (Promega, Madison, WI) and cloned into the pGL3-basic firefly luciferase reporter vector. To accurately map the core region of human CRE1, five deletion mutants R1, R2-5, R3-5, R4-5, R5 were generated by PCR and cloned into the pGL3 basic vector. Further, The R3 regions of human, chimpanzee, gibbon, macaque, marmoset, tree shrew and mouse were amplified and generated by standard restriction enzyme digestion and cloning techniques. All constructs were confirmed by sequencing. The primers used to generate reporter gene constructs are listed in Table S6.

### Generation of Human CCNB1IP1**-**CRE1 and CRE2 KO *SH-SY5Y* cells

For human CCNB1IP1-CRE1 and CRE2 KO using the *SH-SY5Y* cells, single guide RNAs (sgRNAs) were designed using the online CRISPR design tool (Red CottonTM, Guangzhou, China, https://en.rc-crispr.com/). The Human CCNB1IP1 CRE1 and CRE2 regions were selected for CRISPR/Cas9 genome editing. A ranked list of sgRNAs was generated with specificity and efficiency scores. Two pairs of sgRNAs were selected targeting CRE1 and CRE2 respectively, and then the pairs of oligos were annealed and ligated to the sgRNA vector (Ubigene Biosciences, Guangzhou, China). The sgRNA plasmids containing each target sgRNA were transfected into *SH-SY5Y* cells with Lipofectamine 3000 (Thermo Fisher Scientific). 24-48 hours after the transfection, puromycin was added to screen the cells. After antibiotic selection, a certain number of cells were selected, and the KO efficiency of CRE1 and CRE2 SH-SY5Y pool cells were then validated by PCR. The sgRNAs for CRISPR and primers for validation are shown in Table S7.

### Targeting expression vector cloning

The full-length coding region of human *CCNB1IP1* was synthesized by a gene synthesis company (Beijing Tsingke Biotech Co., Ltd.), PCR-amplified, and separately cloned into the GFP-tagged expression vector pCS2-eGFP-N, HA-tagged expression vector pCS2-HA-N, and IUE expression plasmid CAG-IRES-GFP. All final constructs were confirmed via sequencing using an ABI-3130 automatic sequencer. The oligonucleotide primers used for plasmid construction are listed in Table S8.

### *In Utero* Electroporation

*In Utero* electroporation (IUE) was performed as previously described(*72*). Pregnant CD-1 mice were anesthetized using an animal anaesthesia machine (RWD, R500). The skin of the abdomen was cut open using sterilized scissors to pull out the uterus. Next, the ventricle was located in the brain. Using the glass needle, 1–2 µL plasmid was inserted with 0.1% fast green (Sangon Biotech, A610452) as an indicator. A 3-mm electrode positive pole was placed on the side of the blown plasmid, and the electrodes were connected to an electroporator (BTX, ECM830) with the following parameters: 28–30 V, 50 ms duration, 1s interval, and 5–6 cycles. Warm saline was used to maintain the uterus moist and at a constant temperature. Finally, the uterus was returned, the wound was sterilely sutured, and the mice were moved to a warm place to recover.

In all electroporation experiments, the parameters were kept the same. the concentrations of PCAG-CP1 plasmid and control PCAG-GFP plasmid were maintained at 1500ng/µl, the electroporation voltage was maintained at 27 V. The total volume of plasmid injected during the experiment was stable at 1-1.5µl. To obtain reliable statistical results, we used the total number of GFP cells as the denominator to inconsistent electroporation efficiency.

### Immunofluorescence staining and imaging

Samples were perfused or dissected and fixed in 4% paraformaldehyde fixative for 24– 48 h. The tissue was embedded with 4% agarose, and 30–50 µm sections were cut using a Vibratome (Leica, VT1000S). Citrate (Servicebio, G1202) was used for antigen retrieval during immunostaining. The sections were then washed three times in phosphate-buffered saline (PBS) for 5 min and incubated in blocking buffer containing 5% normal donkey serum (Jackson, 017-000-121), 1% bovine serum albumin (BSA) (Sigma, SRE0096), 0.1% glycine, 0.1% lysine, and 0.3% TritonX-100 in PBS for 1–2 h at room temperature (RT). Next, sections were incubated in primary antibody diluted with blocking buffer overnight at 4°C. The sections were incubated with the appropriate fluorescent secondary antibody for 1–2 h at RT in the dark after washing three times in PBS for 5 min each. Nuclei were labelled with DAPI (1:1000, Invitrogen, 62248). Stained sections were mounted using antifade mounting medium (Vector Labs, H-1000). Finally, images were acquired using a confocal microscope (Leica, TCS SP8) and were processed using ImageJ software (National Institutes of Health, NIH).

Primary antibodies: Rabbit monoclonal anti-Ki67 (1:500, Cell Signaling Technology, 9129S); Phospho-Histone H3 (Ser10) Antibody (1:500, Cell Signaling Technology, 9701S); Chicken Polyclonal anti-GFP (1:500, Invitrogen, A10262); Rabbit Polyclonal anti-Pax6 (1:500, Invitrogen, 42-6600); Rabbit Polyclonal anti-TBR2 / Eomes (1:500, Abcam, ab23345); Rat Monoclonal anti-SOX2(1:500, Invitrogen, 53-9811-82); Rabbit Polyclonal anti-SATB2 (1:300, Affinity Biosciences, DF2962); Rabbit Polyclonal anti-TBR1 (1:300, Affinity Biosciences, DF2396); Chicken Polyclonal anti-NeuN (1:500, Millipore ABN91); Mouse anti-nestin (1:500, BD bioscience, Cat#611658); Guinea pig-anti-Ctip2(1:500, Oasis Biofarm, Cat#OB-PGP012) ; Secondary antibodies: Alexa Fluor 594 AffiniPure Donkey Anti-Rabbit (1:800, Jackson ImmunoResearch Labs, 711-585-152); Alexa Fluor 594 AffiniPure Donkey Anti-Rat (1:800, Jackson ImmunoResearch Labs, 712-585-153); Alexa Fluor 488 AffiniPure Donkey Anti-Chicken (1:800, Jackson ImmunoResearch Labs, 703-545-155); Alexa Fluor 488 AffiniPure Donkey Anti-Rabbit (1:800, Jackson ImmunoResearch Labs, 711-545-152); Alexa Fluor 488 AffiniPure Donkey Anti-Rat (1:800, Jackson ImmunoResearch Labs, 712-545-150). Goat anti-guinea pig IgG 594 (1:1000, Oasis biofarm, Cat#G-GP594)

### Cortical Folding Analysis

Brains were carefully isolated from the skull and fixed with 4%PFA for 24-48 hours. Then continuous vibration slicing was performed along the sagittal or coronal direction. Slices were observed under a fluorescence microscope, and samples with gyrification in each litter were counted. slices with cortical folding were singled out for next immunofluorescence staining, including the upper layer marker Satb2, the deep layer marker Ctip2, and radial glial process marker Nestin. The degree of cortical folding was measured with the gyrification index (GI): a ratio of the total cortical inner surface area to the area of an outer surface that smoothly encloses the cortex(*73*). For radial glial process fiber tracing, the direction of each RGC fiber is simulated with lines manually.

### Real-time quantitative RT-PCR

RNA was isolated from N2A cells transfected with 1.6 µg CAG-IRES-GFP empty vector and 1.6 µg CAG-IRES-CCNB1IP1-GFP plasmid to confirm *CCNB1IP1* overexpression. These RNAs were reverse transcribed with oligo-dT(20) primers and amplified with real-time PCR primers using an Applied Biosystems (Waltham, MA, USA) 7500 Real-Time PCR System. The Ct values for each gene amplification were normalized by subtracting the Ct value of the reference gene GAPDH or 18sRNA. Normalized gene expression values are expressed as the relative quantity of *CCNB1IP1* mRNA. The oligonucleotide primers used for qRT-PCR amplification are listed in Table S9.

### Western blotting

Tissues or cells were lysed in RIPA buffer (Beyotime, P0013B) supplemented with a protease inhibitor mixture (Roche, Basel, Switzerland) for 30 min at 4°C. The homogenates were then centrifuged at 13000×*g* for 15 min at 4°C to discard insoluble cell debris. Samples were separated via 10% sodium dodecyl sulphate polyacrylamide gel electrophoresis (SDS-PAGE), transferred onto PVDF membranes, blocked with 5% BSA, and incubated with primary antibodies overnight at 4°C. The following primary antibodies were used for immunoblotting: FLAG (1:5000, Millipore, F1804), HA (1:5000, Cell Signaling Technology, C29F4), CCNP1IP1 (1:1000, Affinity, DF4016), and α-Tubulin (1:5000, Proteintech, 6603-1-Ig). Horseradish peroxidase-labeled anti-rabbit (Cell Signaling Technology, 7074), anti-mouse (Cell Signaling Technology, 7076) or anti-goat (Santa Cruz, sc-2354) secondary antibodies were used. Chemiluminescence was detected using a SuperSignal West Pico PLUS kit (Thermo Fisher Scientific, 34579) according to the manufacturer’s instructions.

Total protein concentration in the brain homogenates or cell lysates were measured using a bicinchoninic acid assay system (Thermo Fisher Scientific). Protein lysates were boiled in SDS loading buffer, and equivalent protein quantities (40–100 g) were resolved using SDS-PAGE and western blotting. Blots were probed with primary antibodies, and immunoreactive bands were quantified using ImageJ software (NIH).

### Cell cycle analysis

The vectors pCS2-GFP and pCS2-GFP-CCNB1IP1 were transfected into N2A cells in 6-well plates using Lipofectamine 3000 and collected 48 h after transfection. Harvested cells were washed twice in PBS, fixed in ice-cold 70% ethanol overnight and mixed on a reciprocating shaker. After 70% ethanol removal through centrifugation (1000 rpm), the cells were washed twice with cold PBS and incubated in 50 μg/mL propidium iodide (Sigma, P4170) with 0.2% TritonX-100 and 100 μg/mL RNaseA (Thermo Fisher, EN0531) for 30 min at 4℃ in the dark. Samples were run on the Becton-Dickinson LSRFortessa flow cytometer (Franklin Lakes, NJ, USA), and data were analyzed using FlowJo software (FlowJo, LLC, Ashland, OR, USA).

### FUCCIO analysis

Neuro2A cells were seeded at low density in a confocal dish (Servicebio, WG801001) with an outside diameter of 35 mm and an inside diameter of 20 mm to determine the duration of the G1 phase. They were then transfected with lipofectamine using two plasmids: Fucci-G1 Orange and control (pCS2-GFP) or pCS2-CCNB1IP1-GFP. Cells were imaged every 2–3 h for 2–4 days at 10× magnification using a high-resolution microscopy (Zeiss, LSM880). A live cell culture system was attached at 5% CO_2_ and 37℃. The number of cells expressing Fucci G1 Orange with GFP was quantified at each time point. We also calculated the proportion and doubling time of cells expressing the G1 marker with pCS2-GFP or pCS2-CCNB1IP1-GFP.

### Behavioural experiments

#### Open-field test

The open-field test(*67*) was used to measure locomotor activity and anxiety-like behavior of 3-month-old male mice. Mice were gently placed in a polyvinyl chloride box (42×42×42 cm) and allowed to explore freely for 5 min. Each mouse was tested in triplicate. The box was thoroughly cleaned after each experiment. The total distance traveled, entries into the center zone, time spent in the center zone, and velocity in both areas were monitored and analysed using Smart v3.0 software (Panlab, Holliston, MA, USA).

#### Novel object recognition (NOR) test

The NOR test is designed to assess short- and long-term recognition memory in mice, and relies on their innate preference for novelty(*74*). The mice roamed freely in the box for 5 min to adapt to the environment. Next, two identical objects were placed and the mice were allowed to explore freely for 10 min. After 24 h, one object was replaced with a novel object. The mice were then allowed to freely explore for 5 min. Exploratory behavior involved touching an object with the mouth or nose and being close to the object (within 2–3 cm).

#### Three-chamber social test

The three-chamber social test was used to assess social deficits and recognition in rodents(*69*). Here, the test apparatus consisted of three interconnected 20×40.5×22 cm chambers. After 5 min habituation to the empty chambers with two empty cages, a same-sex Stranger 1 mouse was placed in Cage 1 and allowed to freely explore for 10 min. Then, another same-sex Stranger 2 mouse was placed in Cage 2. The mice were allowed to freely explore for 10 min. The sniffing time, number of entries, and travel distance into each cage and chamber were then monitored.

#### Morris water maze

The Morris water maze was used to evaluate spatial learning and memory in the rodents(*70*). Here, the water temperature was 18°C -22°C. Mice were trained three times daily for 5–7 days in three random starting quadrants. Each training session lasted for 1 min, and the interval between training sessions was >10 min/d. If the mice did not find the platform within 1 min, they were guided to the platform and allowed to remain there for 15 s. The learning curve was plotted as the number of training days increased. On the last day, the platform in the water maze was removed, and the mice were placed in the quadrant farthest from the platform and allowed to swim freely in the water maze for 1 min. The distance and time spent in each quadrant, escape latency, and escape velocity were recorded.

### Comparison the CCNB1IP1 protein sequence among representative mammalian species

The CCNB1IP1 protein sequences of human, non-human primates and representative vertebrate species were downloaded from the Ensembl database (https://asia.ensembl.org/index.html). Orthologous sequences were aligned using ClustalW (BioEdit software. Tom Hall Ibis Bioscience Carlsbad, CA, USA) and sequence similarity was computed with sequence identify matrix in BioEdit.

### CCNB1IP1 RNA Sequencing read mapping

Illumina reads were processed by trimming adapter sequences using Trim Galore (https://www.bioinformatics.babraham.ac.uk/projects/trim_galore/) and mapped to respective mouse reference genome using HISAT2 2.1.0(*75*). Gene expression was quantified from the mapped reads using HTSeq-count(*76*), which obtained integer counts of mapped reads per gene. Cufflinks software was used to obtain Fragments Per Kilobase Million (FPKM) expression values by automatically estimating library size distributions and sequence composition bias correction(*77*).

### Differentially expressed genes analysis

Differentially expressed genes (DEGs) were identified based on integer count data using the DESeq2 R package(*78*), which determines DE by modeling count data using a negative binomial distribution. Size factors were calculated based on the total number of reads in the different samples. Then, a dispersion parameter was determined for each gene to account for the biological variation between samples. Finally, a negative binomial distribution was used to fit the counts of each gene. *P*-values were calculated using the Wald test, and adjusted for multiple testing with the Benjamini-Hochberg procedure, controlling for the false discovery rate.

### Chinese tree shrew RNA-seq read mapping

Illumina reads were processed by trimming adapter sequences and mapped to the Chinese tree shrew genome TS_3.0 annotation(*34*) using HISAT 2.1.0. Gene expression was quantified from the mapped reads using HTSeq-count(*76*), which obtained integer counts of mapped reads per gene. Cufflinks v2.2.1(*77*) was used to obtain the FPKM expression values. Finally, DEGs were identified based on integer count data using the DESeq2 R package(*78*).

### Human, macaque, and mouse fetal neocortex RNA-seq read mapping

Human and mouse neocortex lamina RNA-seq raw data were downloaded from the Gene Expression Omnibus (GEO) database (www.ncbi.nlm.nih.gov/geoGSE38805). The reads were subsequently processed using FastQC and mapped to the human and mouse indexed reference genomes using HISAT2 2.1.0(*75*). Gene expression from the mapped reads was quantified using HTSeq-count(*76*). Cufflinks software was used to obtain FPKM expression values(*77*). Raw counts and FPKM values for the macaque neocortical lamina were obtained from our previous studies(*43*). DEGs were identified based on integer count data using DESeq2.

### Gene ontology (GO) analysis

We investigated whether DEGs were enriched for biological processes and molecular functions in specific GO terms using the ToppGene Suite(*79*). *P*-values were adjusted for multiple testing using the Benjamini-Hochberg correction (adjusted *p* < 0.05).

### Cluster analysis of gene expression data

The *regularized log* (*rlog*) function transforms original count data to the log2 scale by fitting a model with a term for each sample and a prior distribution on the coefficients estimated from the data. The pheatmap R package was used to perform hierarchical clustering analyses after obtaining the rlog values.

### Comparative analysis of *CCNB1IP1* expression

Both human and macaque *CCNB1IP1* expression values in microarray format(*46, 47*) of each cortical layer at prenatal stages were extracted and reanalyzed from Allen Brain Map (https://portal.brain-map.org/).

The *CCNB1IP1* expression values, TPM, or normalized counts were extracted and reanalyzed from published studies as follows: human and mouse RGC expression data, GSE65000 (*3*); human and gorilla organoid data, GSE153076 (*44*); human, chimpanzee, and bonobo iPS-induced NPCs expression data, GSE124076(*45*); and human, chimpanzee, gorilla, and gibbon eight brain regions expression data, GSE100796(*54*).

Data on the cis- and trans-regulatory effects of genes in human and chimpanzee genes, including *CCNB1IP1*, were downloaded from a published human and chimpanzee-fused organoid study (*56*). The cis-effect was the hybrid log2 (human/chimp) expression, and the trans-effect was the parental log2 (human/chimp) expression minus the hybrid log2 (human/chimp) expression.

### Single-cell data analysis

Human, chimpanzee, gorilla, macaque, and mouse gene expression data from different studies were downloaded from the GEO database. These included human fetal cortex single-cell data (GSE104276), human and chimpanzee organoid or human and macaque fetal cortex data (GSE124299), and human GZ single-cell expression data(*31*). The single-cell analysis R package, Seurat(*80*), was used to read and analyze the feature barcode matrix. The Seurat RunUMAP function was used to perform nonlinear dimension reduction. To perform a comparative analysis between humans, chimpanzees and macaques, we used the FindIntegrationAnchors function to identify anchors that represent cells sharing similar biological states based on canonical correlation analysis(*81*). We then ran integrated analyses on all cells following the step-by-step procedure provided with the Seurat package.

### Data availability

All sequencing data were deposited at the NCBI Gene Expression Omnibus (GEO) under accession number GSE247966 (NCBI GEO).

### Statistical analysis

The *Student’s* t test was used to determine the significant difference by comparing means between two groups of data. Paired data were from the same group but under different conditions. Therefore, in the WB, Luciferase reporter assay, and IUE experiments, two group comparison statistical analysis was performed using an unpaired or paired two-tailed *Student’s t-test*. The CCNB1IP1 control and cKI intersectional comparison was also performed using a paired two-tailed *Student’s t-test*. More than two-group comparisons, we performed One-way ANOVA with post-hoc test to determine significance. A *p-value* of <0.05 was considered statistically significant.

## ACKNOWLEDGMENTS

We thank the members of the Shi Laboratory for their discussions and comments on the manuscript. The FUCCI plasmid was kindly provided by Prof. Guangdun Peng from Guangzhou Institutes of Biomedicine and Health, CAS after signing the MTA. We would like to thank the Core Technology Facility of the Kunming Institute of Zoology, CAS for providing flow cytometry analysis and sorting. We are grateful to Guolan Ma and Shuangjuan Yang for providing technical support.

## FUNDING

L.S. is supported by the Pioneer Hundred Talents Program of the Chinese Academy of Sciences and the Yunnan Revitalization Talent Support Program Young Talent Project. This study was supported by grants from the National Key Research and Development Program of China (2021YFF0702700) to L.S. and BY.M, National Natural Science Foundation of China (32170630), Yunnan Applied Basic Research Projects (202201AS070043 and 202401AS070072), Science and Technology Major Project of Science and Technology of Department of Yunnan Province (202102AA100057), Spring City Project from Kunming Science and Technology Bureau (2022SCP007) to L.S., The joint special project for basic research between the Yunnan Provincial Science and Technology Department and Kunming Medical University (202101AY070001-269) to K.X. NHGRI (HG011641), and National Science Foundation (EF-2204761) to S.V.Y.

## AUTHOR CONTRIBUTIONS

L.S. conceived and designed the study; T.H., YL.T., YF.K., X.K., and JH.W. performed molecular biology and mouse experiments; XL.S. and QF.W. assisted in human fetal cortex samples collection; BY.M. assisted in protein experiments; PC.M. performed protein experiments; L.S. performed bioinformatics analysis; S.V.Y assisted with statistical analyses; L.S., T.H., and YF.K. prepared the figures. L.S., and S.V.Y wrote the manuscript with inputs from other authors.

## DECLARATION OF INTEREST

The authors declare no competing interest.

---

**Fig. S1.**
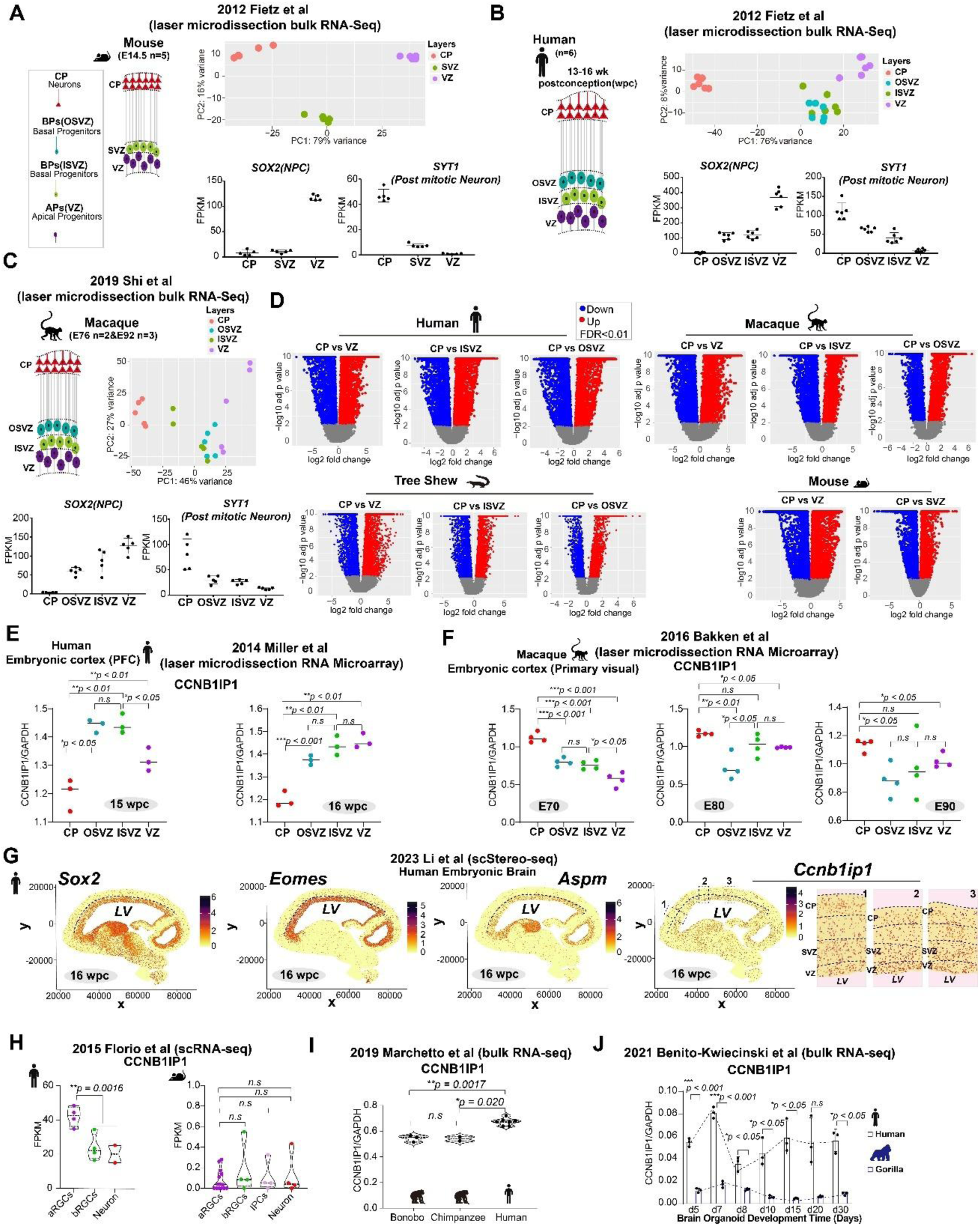
Transcriptome analysis of human, macaque, Chinese tree shrew, and mouse cortex laminae. (**A-C**) PCA maps of mice, humans and macaques based on the expression levels of all genes. Bottom panel: cell marker analysis with *SOX2* for neural progenitor cells and *SYT1* for neurons. (**D**) Volcano plots demonstrating pairwise comparisons of gene expression between the laminae of humans, macaques, Chinese tree shrews, and mice. (**E**) Scatter plots showing pairwise comparisons of CCNB1I1P expression between the laminae of the human fetal cortex at 15wpc and 16wpc respectively (mean, two-tailed unpaired Student’s *t*-test). (**F**) Scatter plots showing pairwise comparisons of CCNB1I1P expression between the laminae of monkey fetal cortex at embryonic 70, 80 and 90 days (mean, two-tailed unpaired Student’s *t*-test). (**G**) Spatial visualization of SOX2, EOMES, ASPM and CCNB1I1P expression in the human fetal cortex at 16wpc. The spatial visualization data were downloaded from brainAtlas (http://donglab.life/brainAtlas.html). (**H**) Violin plot showing *CCNB1IP1* expression in human and mouse aRGC, bRGC, and neurons. (**I**) Violin plot showing *CCNB1IP1* expression in induced pluripotent stem cell (iPS)-derived NPCs among humans, chimpanzees, and bonobos. (**J**) Bar plot showing *CCNB1IP1* expression during human and gorilla brain organoid development (mean, two-tailed unpaired Student’s *t*-test).

**Fig. S2.**
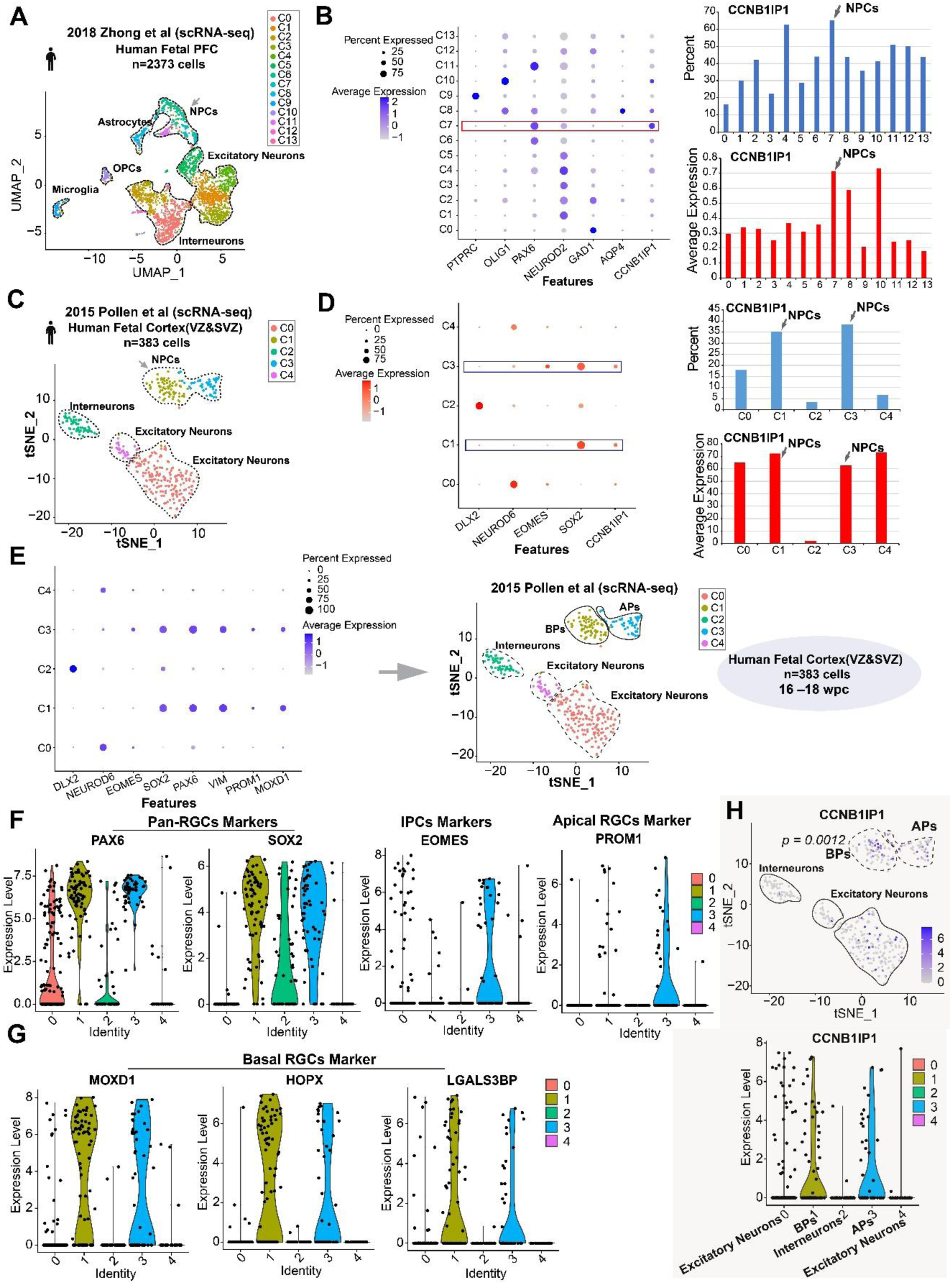
Identification of human *CCNB1IP1* high expression in NPCs, in which, BPs CCNB1IP1 expression is higher than APs. (**A**) Uniform manifold approximation and projection (UMAP) plot showing cell types identified in the human fetal prefrontal cortex (PFC). (**B**) Expression intensity and percentage of *CCNB1IP1* in the identified cell types. (**C**) A t-distributed stochastic neighbour embedding (t-SNE) plot showing cell types identified from the dissected germinal zones of the human fetal cortex. (**D**) Expression intensity and percentage of *CCNB1IP1* in the identified cell types. (**E**) (left panel) Expression intensity and percentage of marker genes in identified cell types; (right panel) a t-SNE plot showing that NPCs cluster3 belongs to APs mixed with EOMES expression and NPCs cluster1 belongs to BPs at human 16-18wpc. (**F**) Violin plots showing the expression of pan-RGCs markers *PAX6*, *SOX2*, IPC marker *EOMES* and apical RGCs marker *PROM1*. (**G**) Violin plots showing the basal RGCs markers *MOXD1*, *HOPX*, and *LGALS3BP* expression pattern. (**H**) Feature and violin plots demonstrating that *CCNB1IP1* expression is higher in BPs than APs at the peak stage of human neurogenesis.

**Fig. S3.**
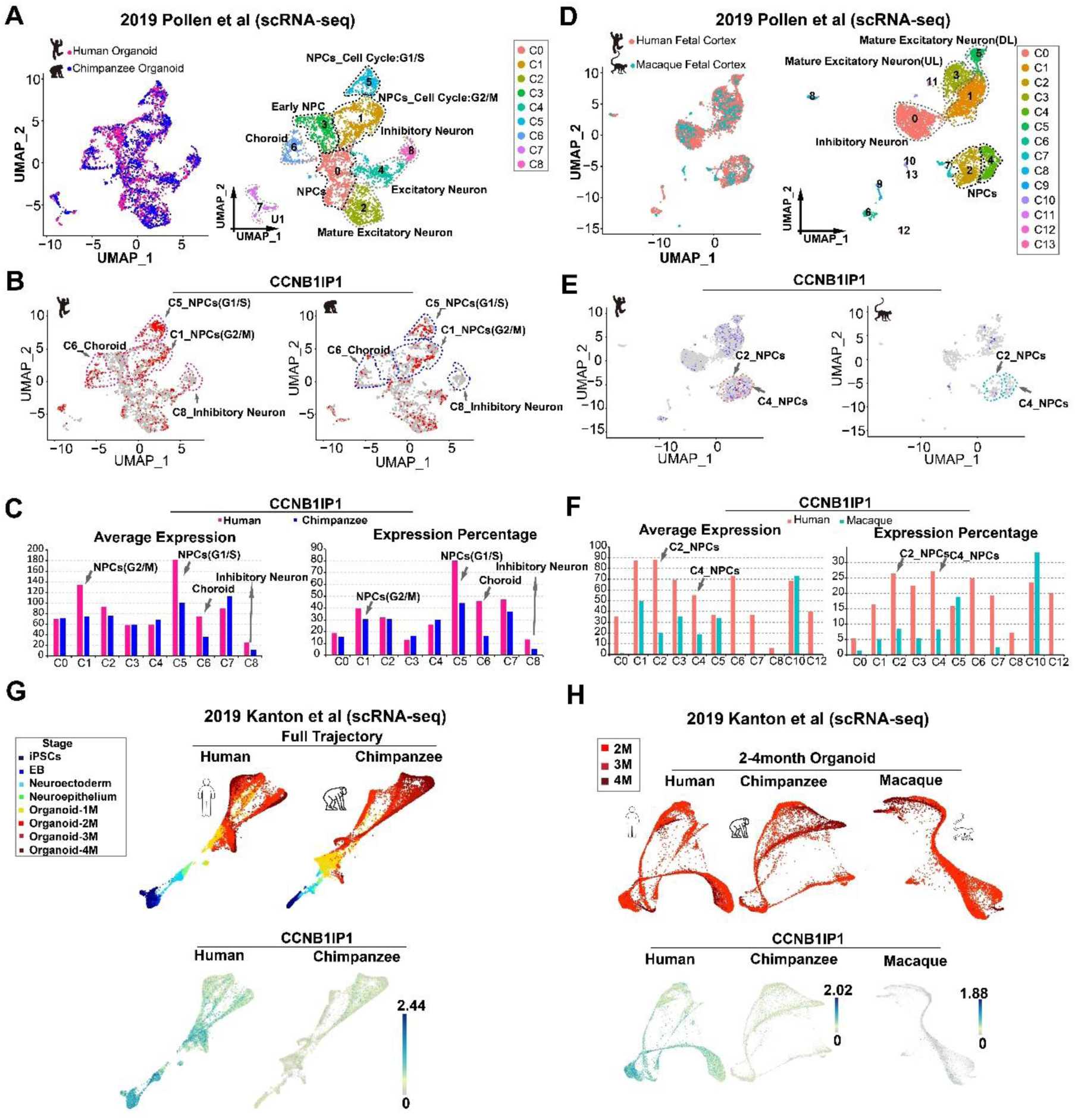
Human *CCNB1IP1* expression is higher than both chimpanzee and macaque in NPCs. (**A**) A UMAP plot showing cell types identified in human and chimpanzee organoids (Pollen et al., 2019). (**B**) UMAP plots showing *CCNB1IP1* expression in human and chimpanzee organoids. (**C**) Expression intensity and percentage of *CCNB1IP1* in the identified cell types. Arrows indicate identified cell types showing higher expression of *CCNB1IP1* in humans. (**D**) A UMAP plot showing cell types identified from human and macaque fetal cortices (Pollen et al., 2019). (**E**) UMAP plots showing *CCNB1IP1* expression in human and macaque fetal cortices. (**F**) Expression intensity and percentage of *CCNB1IP1* in the identified cell types. Arrows indicate identified cell types showing higher *CCNB1IP1* expression in humans. * Clusters 9, 11, and 3 were excluded from the analysis, owing to zero or one cell in the macaque dataset. (**G-H**) *CCNB1IP1* expression in human, chimpanzee, and macaque organoids, indicating high human *CCNB1IP1* expression during organoid development. *Single-cell expression data were downloaded from (https://bioinf.eva.mpg.de/shiny/sample-apps/scApeX/).

**Fig. S4.**
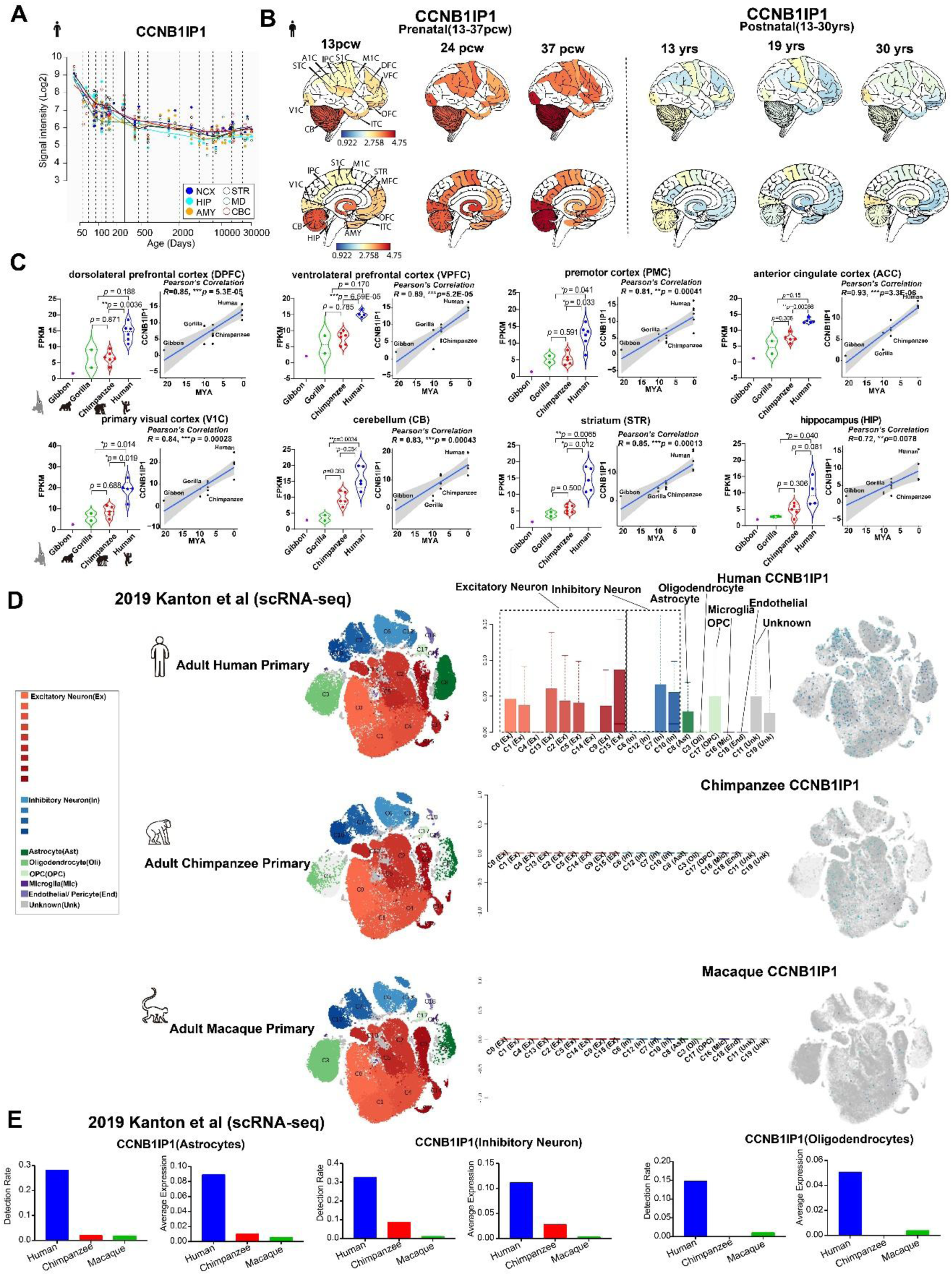
Expression analysis of *CCNB1IP1* at postnatal stages. (**A**) Changes in *CCNB1IP1* expression in brain regions, including the neocortex (NCX), hippocampus (Hip), amygdala (AMY), striatum (STR), mediodorsal nucleus of the thalamus (MD), and cerebellar cortex (CBC) during human brain development. The data were downloaded from the Human Brain Transcriptome Database (http://hbatlas.org/pages/hbtd). (**B**) Spatial *CCNB1IP1* expression changes at the prenatal and postnatal stages. *RNA sequencing data were downloaded from the Brain Span Atlas of the Developing Human Brain at the Allen Institute. (**C**) Violin plots and Pearson’s correlation analysis of *CCNB1IP1* expression in eight brain regions of humans, chimpanzees, gorillas, and gibbons. (**D-E**) *CCNB1IP1* expression in human, chimpanzee, and macaque primary brains. *Single-cell expression data were downloaded from (https://bioinf.eva.mpg.de/shiny/sample-apps/scApeX/).

**Fig. S5.**
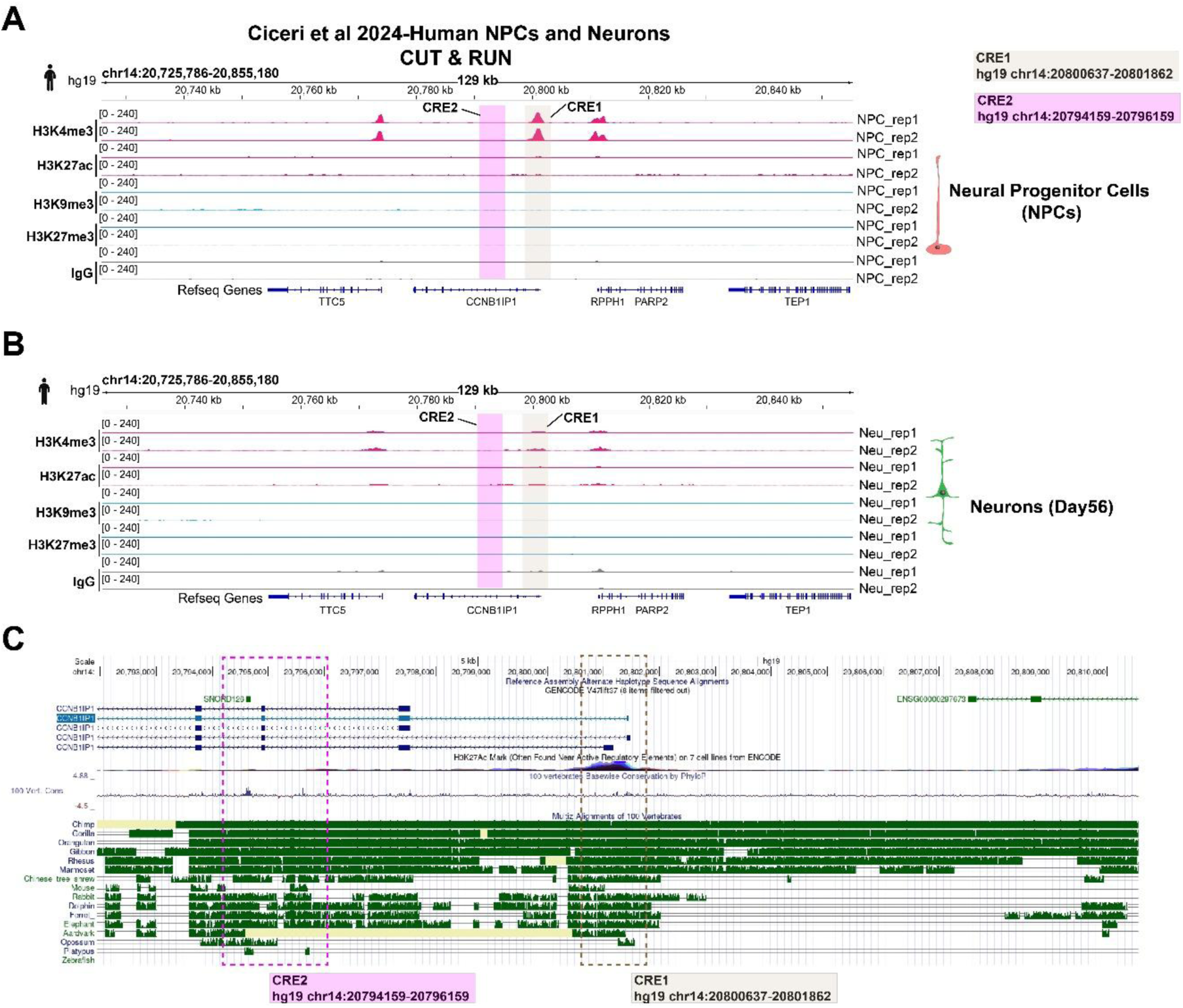
Histone modification and sequence conservation analysis of the *CCNB1IP1* regulatory elements CRE1 and CRE2. (**A-B**) Representative tracks of H3K4me3, H3K27ac, H3K9me3, and H3K27me3 at the CCNB1IP1 genomic loci by analyzing published CUT & RUN data in human NPCs and differentiated neurons. *CRE1 and CRE2 masked with square in brown and pink color respectively. (**C**) Screenshot of CCNB1IP1 genomic loci with multi-species comparisons (green color) using UCSC genome browser. * CRE1 and CRE2 are highlighted in brown and pink square respectively.

**Fig. S6.**
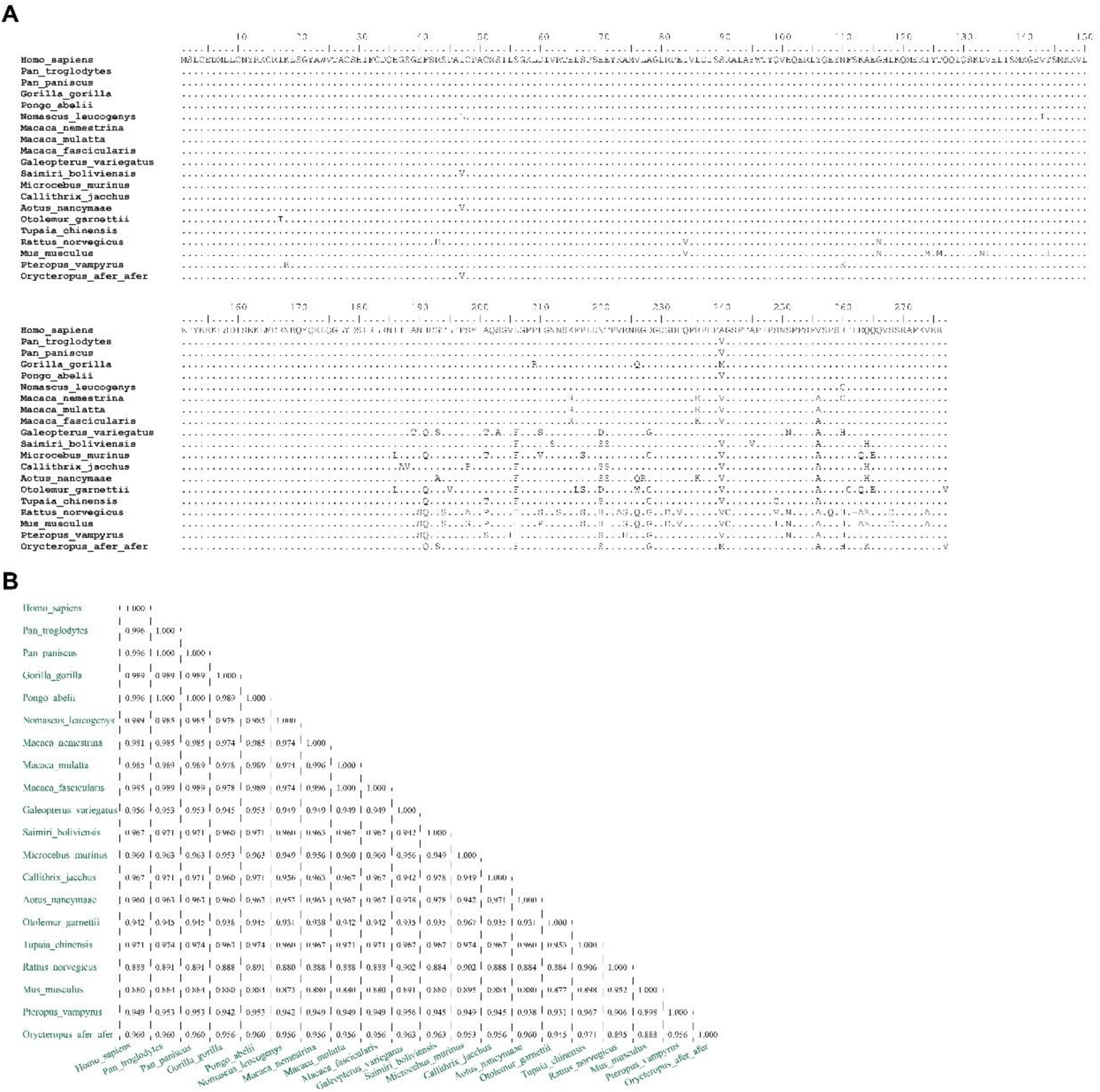
Multi-species comparison analysis of CCNB1IP1 protein sequence. (**A**) Amino acid alignments of the CCNB1IP1 protein sequences of 20 mammalian species. (**B**) Sequence identity matrix results from pairwise comparison of the CCNB1IP1 protein among 20 mammalian species.

**Fig. S7.**
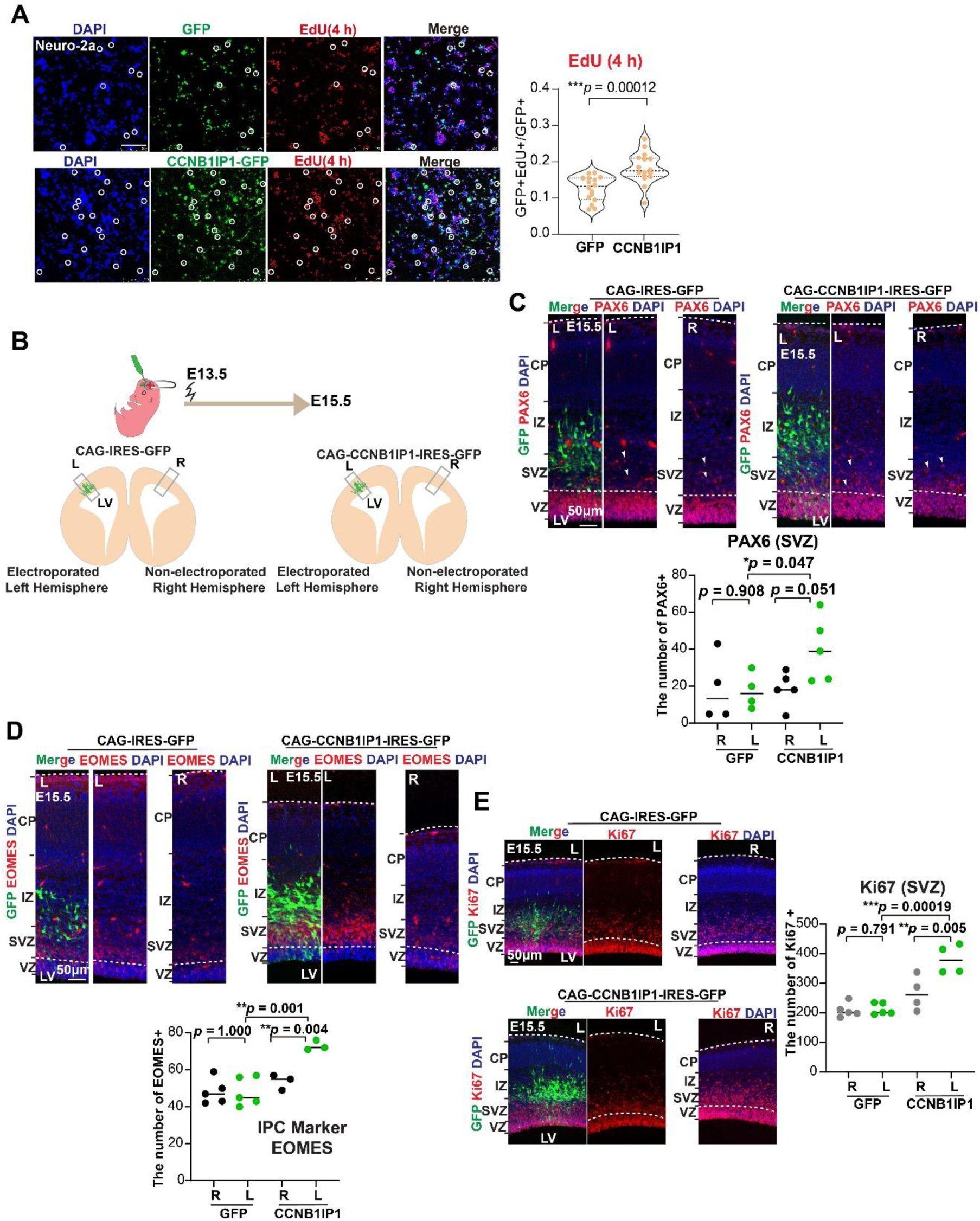
*CCNB1IP1* promotes the proliferation of BPs. (**A**) (Top panel) Representative images of cells transfected with CCNB1IP1-GFP and GFP with EdU pulses for 4 h. (Bottom panel) Violin plots of the percentages of GFP^+^EdU^+^ in GFP^+^ cells. (n=16 vision fields for totally 4 replicates, data represented as mean±SD, \**p* < 0.05, unpaired Student’s *t*-test). (**B**) A schematic diagram of the IUE (E13.5-E15.5) strategy. (**C**) (Top panel) Immunofluorescence images of the APs marker PAX6 at E15.5 in the mouse embryonic cortex. *Arrows indicate PAX6^+^ cells. (Bottom panel) Quantification of PAX6^+^ cells in electroporated and non-electroporated regions, as shown in (a) (n=3-5 embryos from two IUE experiments, data represented as mean, \**p* < 0.05, unpaired Student’s *t*-test). (**D**) (Top panel) Immunofluorescence images of the IPC marker EOMES at E15.5 in the mouse embryonic cortex. (Bottom panel) Quantification of EOMES^+^ cells in electroporated and non-electroporated regions, as indicated in (A), (n=3-5 embryos from two IUE experiments, data represented as mean± SD, \**p* < 0.05, \*\**p* < 0.001, unpaired Student’s *t*-test). (**E**) (Top panel) Immunofluorescence of cell cycle marker Ki67 after electroporation at E13.5. (Bottom panel) quantification of Ki67^+^ cells at electroporated and non-electroporated regions (n=4-5 embryos from two IUE experiments, data represented as mean±SD, \*\**p* < 0.001, \*\*\**p* < 0.001, unpaired Student’s *t*-test).

**Fig. S8.**
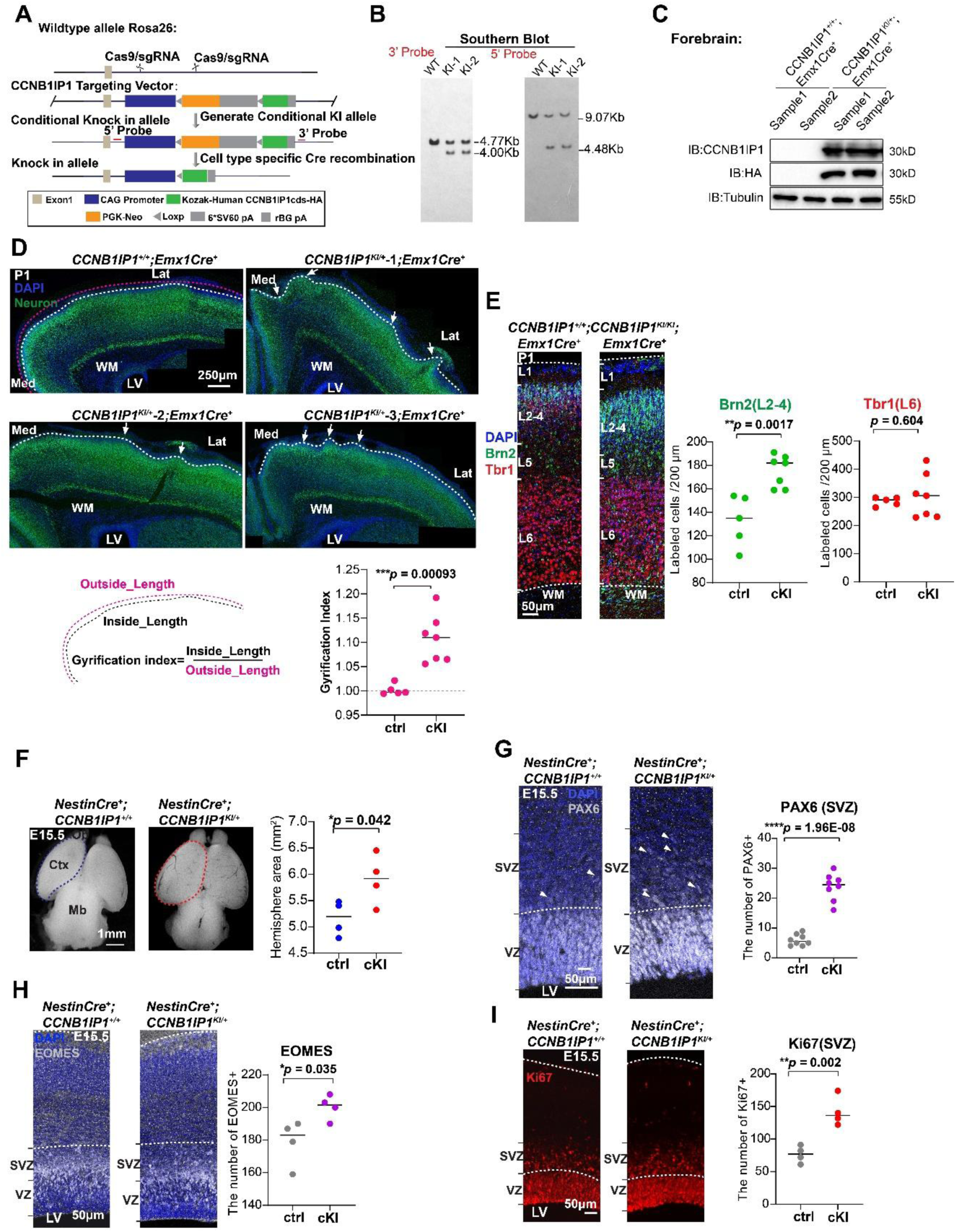
Generation of human *CCNB1IP1* conditional knock-in mice and cortical folding analysis in cKI. (**A**) Schematic representation of the generation of human *CCNB1IP1* conditional knock-in mice at the Rosa26 locus using CRISPR-Cas9 technology. (**B**) A successful knock-in *CCNB1IP1* allele was verified through Southern blotting using 5′ and 3′ probes surrounding the targeting cassette. (**C**) Western blotting analysis of *Emx1-Cre;CCNB1IP1*^KI/+^ cKI mouse cortex, showing high *CCNB1IP1* expression compared to controls. (**D**) (Top panel) Representative images of cortical folding (arrows) in cKI*^Emx1-Cre^* mice at P1; (Bottom panel) Schematic representation and quantification of the degree of cortical folding in cKI*^Emx1-Cre^* mice. Each data point represents a sample, 5 controls and 7 cKI from two nests. (**E**) Immunostaining and quantification of the upper layer marker Brn2 (green) and deeper layer marker Tbr1 (red) in cKI*^Emx1-Cre^* mice, indicating significantly increased upper layer neurons. Each data point represents a sample. 5 controls and 7 cKI from two nests. (**F**) A dorsal view of the control and cKI*^Nestin-cre^* embryonic brains at E15.5 (left panel) and quantification results showing significant increase of the cortical area in cKI*^Nestin-cre^* mice (right panel). Each data point represents a sample, 4 controls and 4 cKI from one nest. (**G-I**) Representative images and quantification of PAX6^+^, EOMES^+^, and Ki67^+^ cells in cKI*^Nestin-cre^* mice. Each data point represents a sample. All statistical data are presented as the mean or mean, \**p* < 0.05, \*\**p* < 0.001, \*\*\**p* < 0.001, *n.s*, not significant, as determined using the unpaired Student’s *t*-test.

**Fig. S9.**
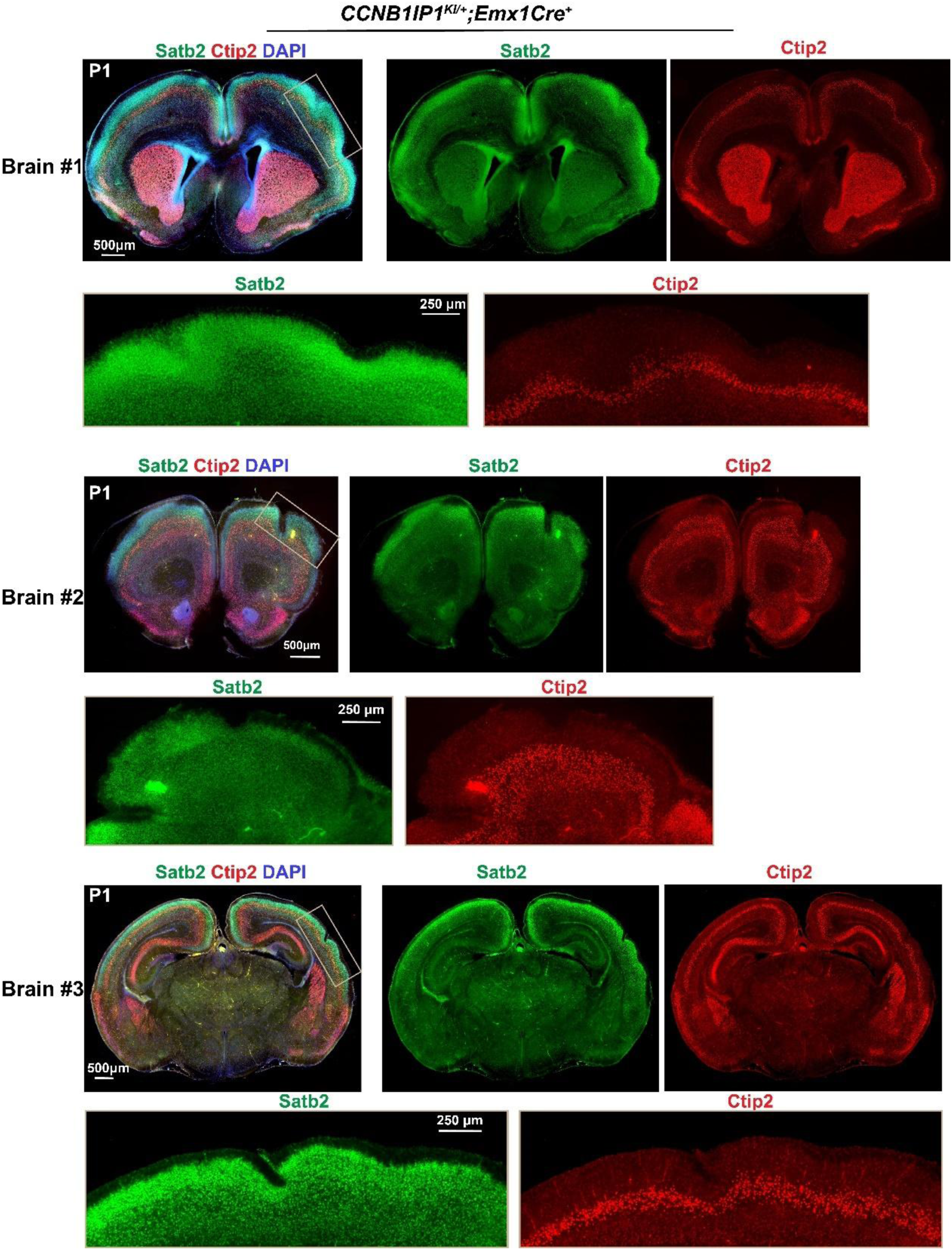
cortical folding analysis of cKI at P1. Representative images of cortical layer marker SATB2 (L2-4) and CTIP2 (L6) in three independent cKI mice collected from different nests at P1. Area with white rectangle to be magnified to show the gyri and adjacent sulci structures in the folding locations.

## Notes

### Competing Interest Statement

The authors have declared no competing interest.

